# Motoneuron-derived Activinβ regulates Drosophila body size and tissue-scaling during larval growth and adult development

**DOI:** 10.1101/661363

**Authors:** Lindsay Moss-Taylor, Ambuj Upadhyay, Xueyang Pan, Myung-Jun Kim, Michael B. O’Connor

## Abstract

Correct scaling of body and organ size is crucial for proper development and survival of all organisms. Perturbations in circulating hormones, including insulins and steroids, are largely responsible for changing body size in response to both genetic and environmental factors. Such perturbations typically produce adults whose organs and appendages scale proportionately with final size. The identity of additional factors that might contribute to scaling of organs and appendages with body size is unknown. Here we report that loss-of-function mutations in Drosophila *Activinβ (Actβ)*, a member of the TGF-β superfamily, lead to production of small larvae/pupae and undersized rare adult escapers. Morphometric measurements of escaper adult appendage size (wings, legs), as well as heads, thoraxes, and abdomens, reveal a disproportional reduction in abdominal size compared to other tissues. Similar size measurements of selected *Actβ* mutant larval tissues demonstrate that somatic muscle size is disproportionately smaller when compared to fat body, salivary glands, prothoracic glands, imaginal discs and brain. We also show that Actβ control of body size is dependent on canonical signaling through the transcription-factor dSmad2 and that it modulates the growth rate, but not feeding behavior, during the third instar period. Tissue and cell-specific knockdown and overexpression studies reveal that motoneuron derived Actβ is essential for regulating proper body size and tissue scaling. These studies suggest that, unlike in vertebrates where Myostatin, and certain other Activin-like factors act as systemic negative regulators of muscle mass, in Drosophila Actβ is a positive regulator of muscle mass that is directly delivered to muscles by motoneurons. We discuss the importance of these findings in coordinating proportional scaling of insect muscle mass to appendage size.

## Introduction

Some members of the animal kingdom, including most species of fish, amphibians, lizards, turtles, and salamanders, undergo indeterminate growth and increase their biomass throughout their lifespan. In contrast, birds, mammals and many insect species exhibit determinate growth whereby ideal body length and weight is fixed upon reaching sexual maturity. This process produces a more limited range of sizes that are characteristic for the species (HARIHARAN *et al*. 2015). In these animals, growth rate can vary during development and is influenced by both intrinsic and extrinsic factors. For example, in humans, at the conclusion of the high pubertal growth period, the long bone growth plates are ossified thereby preventing additional increase in overall skeletal size (KRONENBERG 2003; SHIM 2015). Similar to mammals, holometabolous insects also exhibit determinate growth. In Drosophila, a larva increases its mass 200-fold (70% of which occurs in the last larval instar) before terminating growth at pupariation (CHURCH AND ROBERTSON 1966). During the non-feeding pupal stage, the adult structures differentiate from larval imaginal tissue and there is no net increase in body mass. Thus, the final body size is set by the rate of larval growth and the timing of its termination.

In recent years, numerous studies have centered on elucidating the molecular mechanisms that regulate hormonal activity during larval development in holometabolous insects to better understand how growth rate and duration are controlled (reviewed in (REWITZ *et al*. 2013; BOULAN *et al*. 2015). In Drosophila, growth is largely regulated by the Insulin/IGF Signaling (IIS) and Target of Rapamycin (TOR) pathways, which are themselves regulated by different nutritional inputs. IIS is regulated by systemic sugar concentrations and TOR by circulating amino acid levels. Mutations that attenuate either pathway lead to slower growth rates resulting in diminutive animals with smaller and fewer cells ((CHEN *et al*. 1996; BOHNI *et al*. 1999; OLDHAM *et al*. 2000; RULIFSON *et al*. 2002)). Conversely, activation of either pathway can lead to larger organs and cells if there are adequate nutrients (LEEVERS *et al*. 1996; GOBERDHAN *et al*. 1999; STOCKER *et al*. 2003). Interestingly, systemic manipulation of IIS/TOR pathways typically leads to smaller or larger animals, with proportional effects on organ and appendage size (allometric growth) (SHINGLETON *et al*. 2007; SHINGLETON AND FRANKINO 2013).

While IIS/TOR are central regulators of growth rate in holometabolous insects, the major regulator of growth duration is the steroid hormone 20-hydroxyecdysone (20E) (reviewed in (YAMANAKA *et al*. 2013a). During the final larval stage, a pulse of 20E extinguishes feeding, terminates growth and initiates pupariation. The timing of the 20E pupariation pulse is triggered, in part, by the neuropeptide prothoracicotropic hormone (PTTH) which in Drosophila is produced by the two pairs of neurons in each brain hemisphere that innervate the prothoracic gland (PG) (MCBRAYER *et al*. 2007; SHIMELL *et al*. 2018). PTTH binds to its receptor Torso and stimulates synthesis and secretion of ecdysone from the PG (REWITZ *et al*. 2009; YAMANAKA *et al*. 2013a). PTTH production/release responds to a variety of environmental signals including nutritional status, light, and tissue damage as well as internal signals such as juvenile hormone (JH) to further tune the timing of pupariation (YAMANAKA *et al*. 2013b; DE LOOF *et al*. 2015; SHIMELL *et al*. 2018).

In addition to IIS/TOR signaling and steroid hormones, other signaling pathways have also been identified that affect final body mass and proportion scaling in both vertebrates and invertebrates. In particular, the TGFβ signaling pathway has known roles in controlling cell, tissue, and body size. TGFβ super-family ligands signal by binding to a heterotetrameric complex of type I and type II serine-threonine receptor kinases. Ligand binding triggers type II receptors to phosphorylate type I receptors, thereby activating its kinase (HELDIN AND MOUSTAKAS 2016). In canonical signaling, the activated type I receptor phosphorylates its major substrates, the R-Smads (reviewed in (HATA AND CHEN 2016). Once phosphorylated, R-Smads oligomerize with a co-Smads and translocate to the nucleus where together with other cofactors they regulate gene transcription (review in (HILL 2016). The ligand super-family is broadly divided into two major sub-divisions based on phylogenetic and signaling analysis (KAHLEM AND NEWFELD 2009). These include the TGFβ/Activins, which in vertebrates signal through R-Smads 2/3, while the BMP/GDF-type factors signal through R-Smads 1/5/8 (MACIAS *et al*. 2015).

TGFβ family members contribute to tissue and body size growth by a variety of mechanisms. For instance, in mammalian mammary cells, TGFβ cell-autonomously regulates cell size via mTOR during epithelial-mesenchymal transition (EMT) (LAMOUILLE AND DERYNCK 2007). In addition, BMPs have been shown to control cell proliferation at the long bone growth plate and have been identified by genome-wide association studies as regulating human height (HIRSCHHORN AND LETTRE 2009; WOOD *et al*. 2014). Another particularly stunning example is Myostatin, a circulating Activin-type ligand, whose loss causes skeletal and muscle hypertrophy in vertebrates (MCPHERRON *et al*. 1997; MCPHERRON AND LEE 1997). TGFβ-type factors also affect the body size of invertebrates. For example, in *C. elegans*, a BMP-type ligand *DBL-1*, is secreted from neurons and signals via *small* (*sma*), a worm Smad, in the hypodermis to regulate expression of cuticle genes (TUCK 2014; MADAAN *et al*. 2018).

To further explore how TGFβ ligands influence body size, we investigated the role of Drosophila Actβ in regulating these traits using both loss and gain-of-function studies. In Drosophila, genetic studies as well as phylogenetic analysis suggests that Actβ signals via Baboon (Babo) and Punt, type I and type II receptors respectively, to phosphorylate dSmad2 (reviewed in (UPADHYAY *et al*. 2017). We find that canonical Actβ signaling through dSmad2 regulates adult viability, body size, and tissue scaling. *Actβ* mutants produce small larvae and pupae along with rare adult escapers. Compared to controls, these rare mutant adults exhibit small abdomens while other structures such as the head, thorax, leg, and wing are of relatively normal size. In larvae, muscle size is most profoundly affected while imaginal discs and the larval brain are of normal size. Furthermore, *Actβ* mutants have a slower overall growth rate, but show no defects in food intake. Using tissue specific gain and loss-of-function, we demonstrate that motoneuron derived Actβ is required for proper muscle growth and adult viability. Conversely, hyperactivation of Activin signaling in muscles by overexpression of activated Babo produces a much larger animal with bigger muscles, but smaller imaginal discs. These observations demonstrate that muscle size can be perturbed without having proportional effects on the size of the imaginal tissues. Therefore, we suggest that either muscles and appendages do not typically coordinate their growth or that such coordination requires *Actβ* signaling.

## Results

### Actβ is required for adult viability, normal body size, and correct tissue scaling

Drosophila *Actβ* has been shown to be involved in a diverse group of developmental processes, including neuroblast proliferation, photoreceptor tiling, regulation of Akh signaling, inter-organ regulation of mitochondrial and hemocyte function (TING *et al*. 2007; ZHU *et al* 2008; MAKHIJANI *et al*. 2017; SONG *et al*. 2017a; SONG *et al*. 2017b). However, in none of these studies was the lethal stage or the gross morphological phenotype carefully documented. To examine this issue, we initially characterized mutant phenotypes using the previously reported putative *Actβ^ed80^* null allele (nonsense mutation) (ZHU *et al*. 2008). However, since the *Actβ* locus is on the fourth chromosome, additional recessive background mutations on the *Actβ ^ed80^* chromosome cannot be removed by recombination and therefore could complicate the phenotypic analysis of homozygous *Actβ^ed80^* mutants. To resolve this issue, we generated several independent deletion alleles (*Actβ^4E^, Actβ^10E^* and *Actβ ^4dd^*) in the *w^1118^* background using the CRISPR/Cas9 system (REN *et al*. 2013; SEBO *et al*. 2014). All phenotypes initially described using *Act^ed80^* homozygotes were confirmed using different combinations of transheterozygous alleles to rule out 4^th^ chromosome background effects.

All examined *Actβ* mutant alleles are predominantly late pupal (pharate) stage lethal (Fig. 1A and B). Many of the pharates show limited movement inside the pupal case, but most never eclose. Manual cracking of the operculum allowed a small percentage (∼1%) to escape and produce viable adults that exhibit severe locomotive defects, and held out immobile wings rendering them flightless (Movie 1,2). Despite these behavioral/physical defects, females could mate and produce offspring from wildtype males. *Actβ* mutant males were unable to produce progeny with either mutant females or wildtype females. Whether this is a behavioral issue (i.e. unable to initiate courtship behavior) or a fertility defect was not determined.

**Figure 1.**
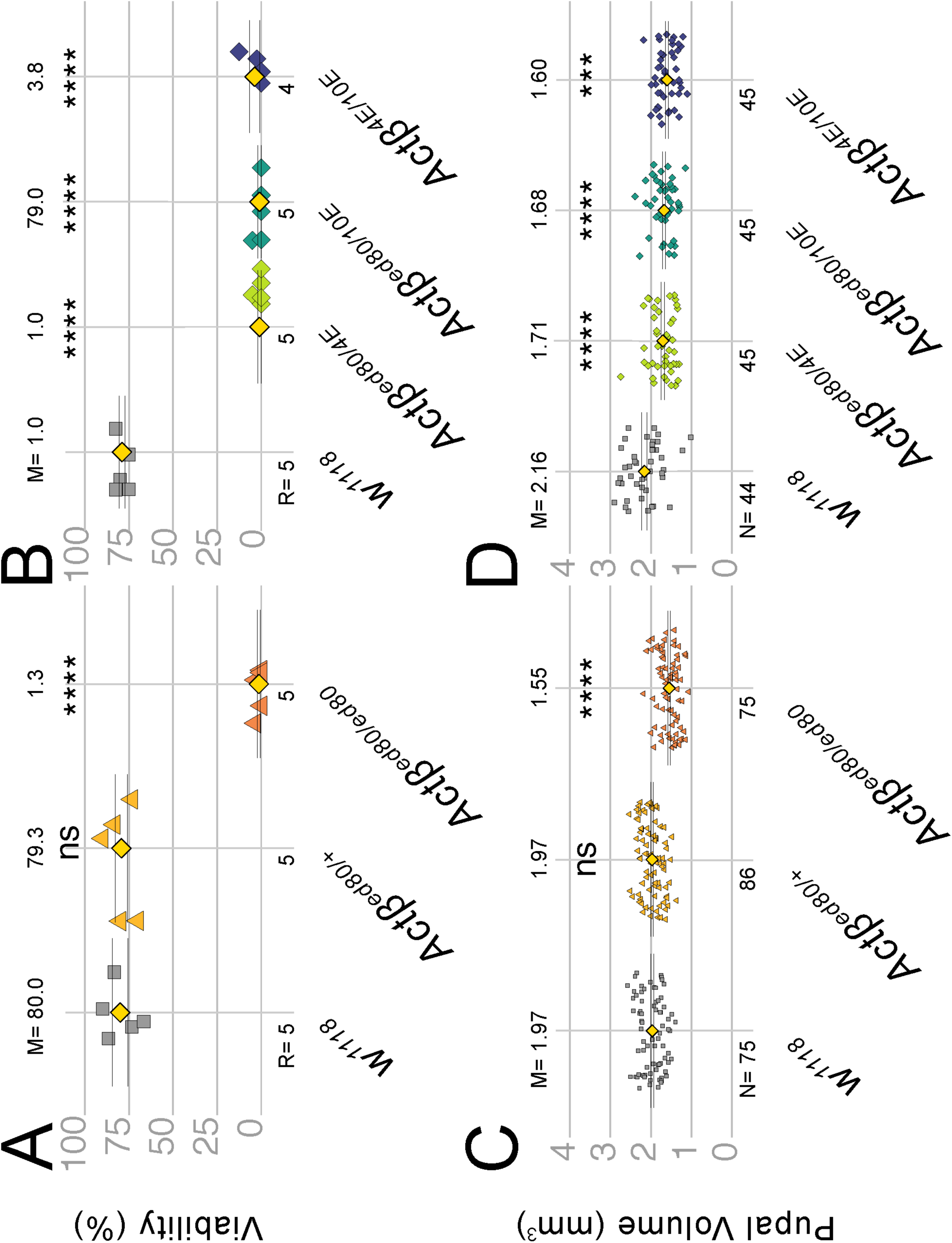
*Actβ* null mutants exhibit a small body size and late pupal lethality. (A,B) Most *Actβ* mutants die as late pharates in the pupal case with between a 1-4% escaper rate. Heterozygotes and *w^1118^* controls exhibit ∼80% viability. (C) Pupal volume of *Actβ^ed80^* (mixed male and female pupa) null mutants (orange triangles, 1.55 mm^3^) are ∼20% smaller than heterozygous individuals (yellow triangle, 1.97 mm^3^) and *w^1118^* controls (grey squares, 1.97mm^3^) (D) Pupal volumes of other A*ctβ* trans-heterozygous mutant combinations show similar decreases in pupal volume. M is the sample mean shown above each data set, N is sample size for pupal volume and R is number of replicates for each genotype (A,B), each replicate consists of 30 - 40 larvae. Means indicated by yellow diamond, ± SEM.

In addition to pharate lethality, *Actβ* mutants exhibit a small body size at all stages of development. *Actβ^ed80^* homozygous pupae (mixed male and female populations) are 21% smaller by volume relative to *w^1118^* or heterozygous pupae (Fig. 1C). Similar to the *Actβ^ed80^* homozygous phenotype, all trans-heterozygous combinations (*Actβ^ed80/4E^*, *Actβ^10E/ed80^*, *Actβ^10E/4E^*) are also significantly smaller (21%, 22%, and 26%, respectively) compared to *w^1118^* control (Fig 1D) indicating that the small pupal size is not caused by secondary mutations on the mutant chromosome. Taken together, these data indicate that *Actβ* is required to produce normal pupal volume and adult viability.

Appendage size is proportionally scaled with body mass in Drosophila (Mirth and Shingleton, 2012). To examine if the adult body components of *Actβ* mutants are proportionally reduced, we collected 1 day old escaper males and females and measured various traits. We found that the *Actβ^ed80^* homozygous male weights are reduced on average 28% compared to the control (Fig. 2A, female 20% not shown). We next measured the abdomen, thorax, and prothoracic leg lengths, along with head projection area and wing surface area of *Actβ* mutant males and controls. Interestingly, the size of some adult structures of *Actβ* mutants are more severely affected than others (Fig. 2B-L). The abdomen length in *Actβ* mutants is reduced by a much greater proportion, −24% (Fig. 2D, F) than any other measured component: head projection area, −8% (Fig. 2B, C), thorax length, −4% (Fig. 2D, E), prothoracic leg length, −2% (Fig. 2G, J), wing area −4% (Fig. 2 H, K). Using the wing trichome density as a proxy, we found no difference in cell size between *Actβ* mutants and the *w^1118^* control (Fig. 2K, I, L), indicating that the minor reduction in wing size is likely caused by a subtle defect in cell proliferation at some time during development.

**Figure 2.**
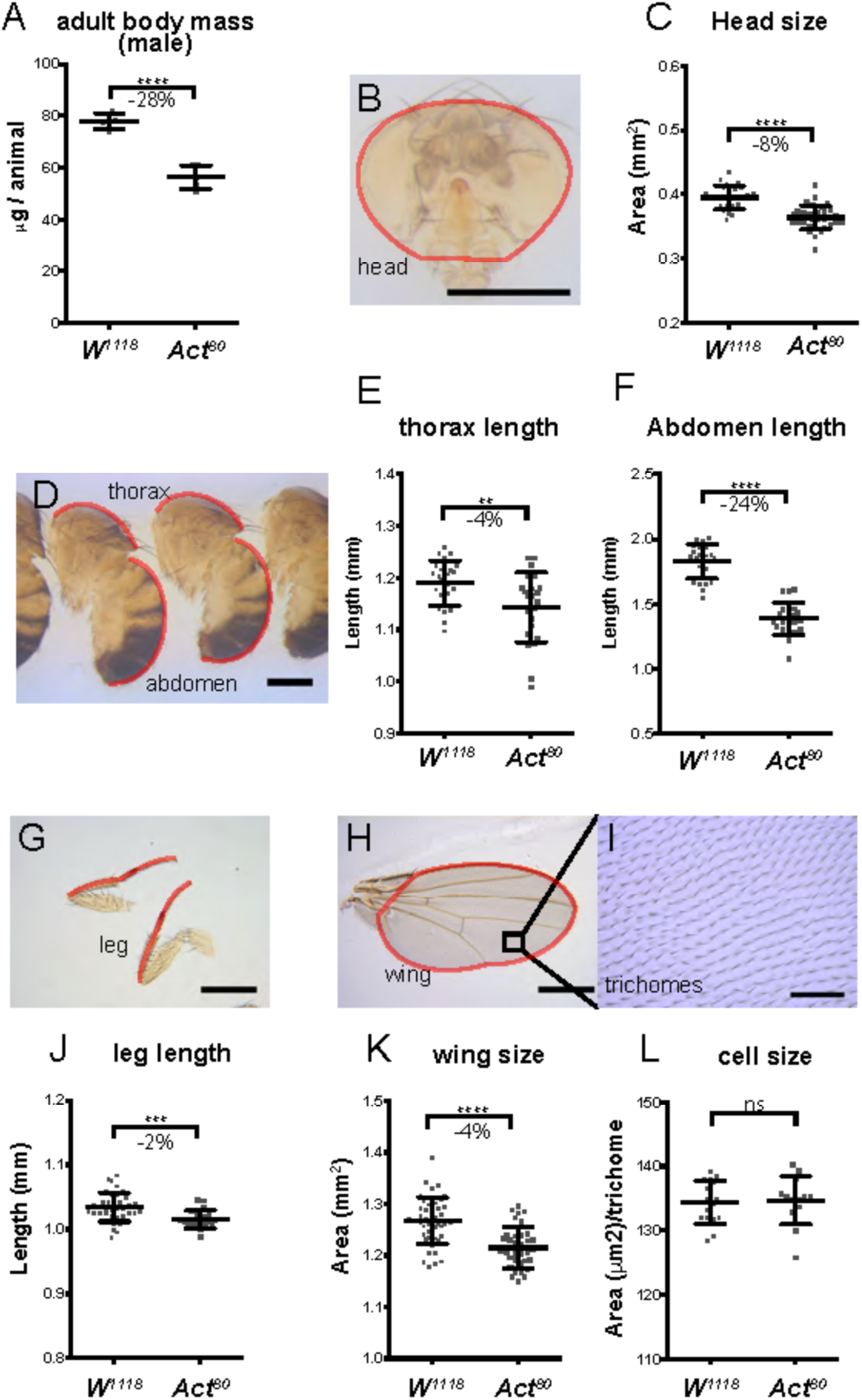
*Actβ* mutant adult escapers have a disproportionately smaller abdomen compared to head, thorax, leg, or wing. (A) *Actβ^ed80^* mutants that eclose as adults weight ∼ 28% less than vs *w^1118^* controls (n = 3-4 groups containing 9-10 individuals). (B-C) Heads of mutant males are ∼8% smaller, (scale 500μm, n>30). (D) Thorax and abdomen (F) are ∼4% and ∼24% smaller respectively, (scale 500μm, n = 23). (G,J) Legs and wings (J,K) of mutants are ∼2% and ∼4% smaller respectively (scale 500μm, n = 22 – 46). (I, L) trichome density in the adult wing shows no difference in cell size (scale 50μm, n = 13 – 16). Means ± S.D. are shown.

### Actβ disproportionately affects larval muscle and certain polyploid tissue sizes

To understand the size discrepancy of adult structures in *Actβ^ed80^* mutants, we examined directly the size of various larval tissues including brain, wing and leg discs, and body-wall muscles, and indirectly the sizes of several polypoid tissues including the fat-body, proventriculus, salivary and prothoracic gland cells using the size of the nucleus as a proxy for cell size.

The most pronounced defect of *Actβ^ed80^* mutant larvae is exhibited by the body wall muscles which in males are reduced by 37% (Fig. 3 A-C), and muscle nuclear size by 53% (Fig. 3D-F). The muscle size reduction is not caused by an earlier myoblast fusion defect since mutant muscles contain the same number of nuclei as wildtype (Fig. S1). In contrast, neither the brain volume (Fig. 3J, K) nor the 2-D projected surface area of the wing, leg, and haltere disc (Fig. 3K-P) were significantly affected. Interestingly, we note that the nucleus size of several other polyploid tissues including the fat body and the prothoracic glands are also significantly reduced, but to a lesser degree than the muscle nuclei (22% for fatbody, Fig. 3FD-I, 37% prothoracic gland vs 53% muscle Fig S2 G-I). Curiously, the nuclear sizes of the cells within the proventriculus remained unchanged (Fig. S2 A-C) and the average salivary gland cell nuclear size actually increased. (Fig. S2 D-F). We conclude that the small pupal volume and reduced escaper weights are primarily due to the disproportionate reduction in muscle size rather than alterations in mitotic tissue growth such as the brain and imaginal discs.

**Figure 3.**
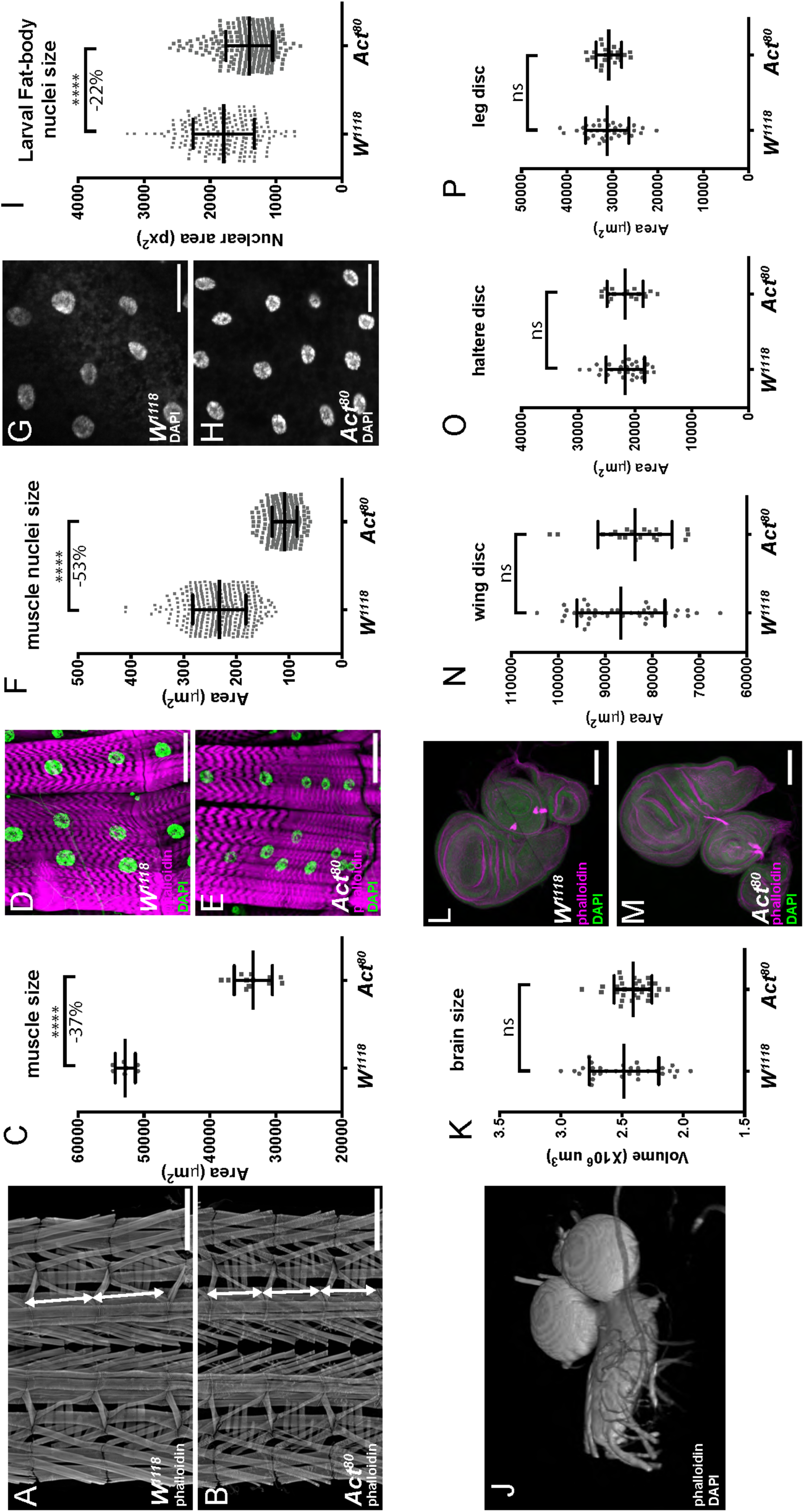
Actβ disproportionately affects the size of larval body wall muscles and fat body nuclei size. Late wandering L3 larvae were dissected and the size various tissues determined. (A,B) Larval fillets were stained with rhodamine-phalloidin and imaged in the muscle plane. Double headed arrows mark the extents of a larval segment photographed at the same magnification (scale bar = 500μm). Note that approximately 3 segments of *Actβ* mutant muscles occupy the same area as 2 wildtype segments. (C) Loss of *Actβ* results in a 37% decrease in the surface area of muscle #6 from the A2 segment compared to control (n = 7 – 12). (D - F) Muscle nuclei (DAPI, green) of *Actβ* mutants are 53% smaller (scale bar =50μm, n > 250) than controls. (G – I) Fat body nuclei (DAPI, gray) of *Actβ* mutants are 22% smaller than control (scale bar = 50μm, n > 200). (J – K) 3D reconstruction of larval brains stained with DAPI and rhodamine-phalloidin, the volume of each brain lobe was measured separately and *Actβ* mutants showed no significant difference of brain size compared to control (n > 30). (L-P) Wing, leg, and haltere imaginal discs (DAPI green, phalloidin magenta) of *Actβ* mutants are the same size as controls, scale 100μm, (n > 20 in each group). Means ± S.D. are shown.

### *Actβ* mutants feed normally but grow slowly

Body size is largely determined by two factors, the duration of growth and the growth rate, or some combination of the two parameters. In addition, a slower growth rate may reflect reduced food intake, diminished absorption of nutrients, or an alteration in metabolic flux. We examined several of these parameters to determine if they were altered in *Actβ* mutants. First, we measured the larval growth rate during the L3 period, when most of the larval growth occurs. At the start of the L3 stage, there is no difference in mass of the mutants versus the controls; however, over the course of 36 hours, a slower rate of mass accumulation becomes apparent such that, at the time when larvae begin to wander, the *Actβ^ed80^* mutants weigh 18% less than *w^1118^* controls (Fig. 4A). This difference in growth rate likely accounts for a large portion of the reduced body size phenotype. To examine whether the diminished growth rate might reflect reduced food intake, we measured feeding rates of foraging early L3 larvae by the mouth-hook contraction assay (WU *et al*. 2003; WU *et al*. 2005). Surprisingly, we found no difference in the head contraction rates of the *Actβ* mutants (Fig. 4B) suggesting that the slow growth rate of these mutants is not likely caused by reduced food intake, but instead may reflect an alteration in nutrient absorbance or dysfunctional metabolic flux.

**Figure 4.**
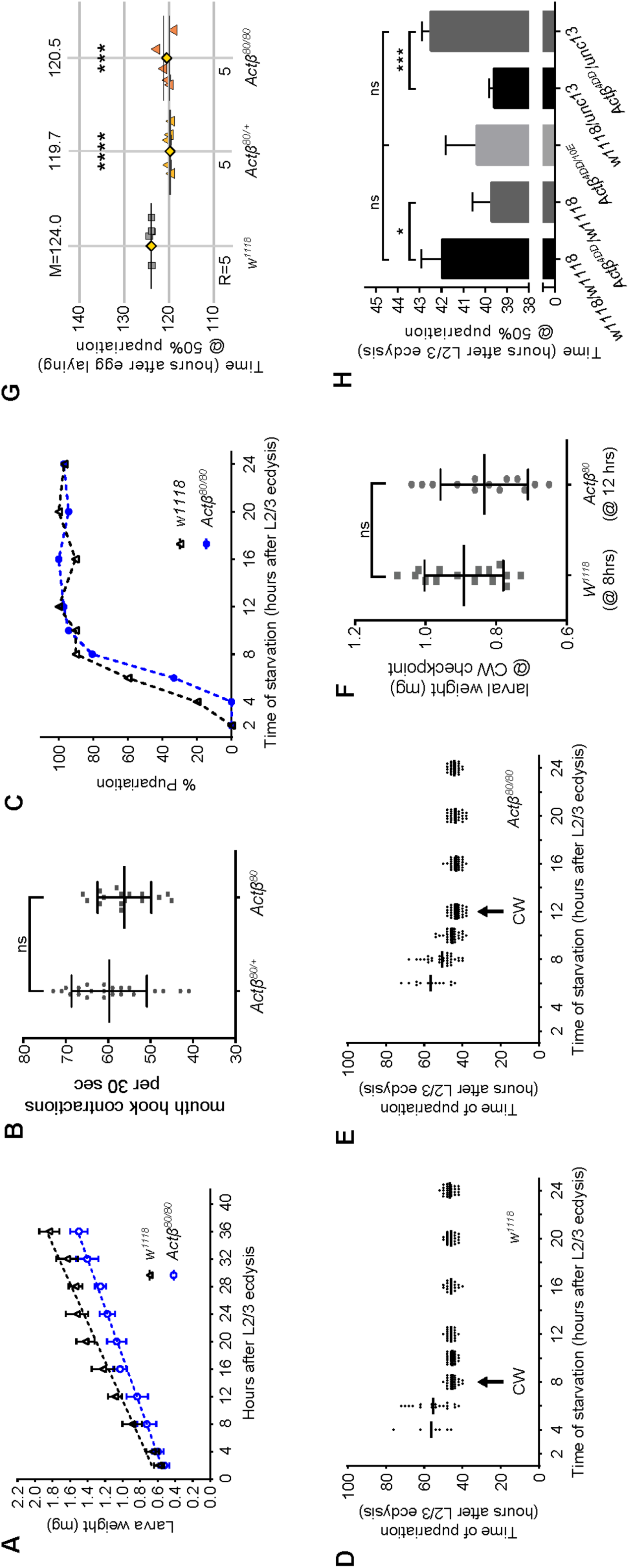
*Actβ* mutant larvae grow slowly but do not exhibit differences in Critical Weight or Developmental timing. (A) Mutant and control larvae were synchronized at L2/L3 ecdysis then larval wet weight was measured at various time intervals. *Actβ* mutants (blue circles/line) weigh the same as *w^1118^* controls (black triangles/line) immediately after L2/3 ecdysis but after 36 hours weigh ∼18% less than controls, (n = 8 – 10 per group). (B) Mouth hook contraction assay of early L3 larvae finds no difference in feeding rates of *Actβ* (squares) mutants vs controls (circles) (n = 16 – 22). (C) *Actβ* mutants are more sensitive to starvation in the early L3 stage than controls (D, E) The critical weight checkpoint was determined by identifying the time at which starvation does not delay pupariation. *Actβ* mutants reach the CW checkpoint 4 hours after controls. (F) Comparing the larval mass when larvae reach their respective CW checkpoints shows that Actβ (at 12 hrs AL2/3) weight the same as controls (at 8 hrs AL2/3). (G) Developmental timing analysis of when 50% of the population have pupariated of *Actβ^80^* mutants and heterozygotes vs controls, shows that both mutants and hets pupariate ∼5 hrs earlier mean ± SEM). (H) Developmental timing analysis of CRISPR alleles and various heterozygous controls. *Actβ^4dd/10E^* mutants do not develop faster than the *w^1118^* control, however both *Actβ^4dd^/w^1118^* and *unc13GFP/w^1118^* hets develop ∼3 hrs faster mean ± SEM than either *w^1118^* or the *Actβ^4dd/10E^* mutant combination. Unless indicated mean ± SD is shown.

Next, to determine whether the small body size might also involve a reduced growth period, we measured the time to pupariation as well as the critical weight (CW), which is a nutritional checkpoint that ensures larvae have enough nutrient stores to produce viable adults (NIJHOUT AND CALLIER 2015). In both the control and *Actβ^ed80^* mutant, starvation after just 2 hours into the L3 stage blocks pupariation (Fig. 4C). In the *Actβ^ed80^* mutant, starvation beginning 4 hours after L2/L3 ecdysis results in delayed pupariation, while starvation 12 hours after L3 ecdysis results in no developmental delay, indicating attainment of CW. The *w^1118^* control achieves CW 8 hours after L2/L3 ecdysis (Fig. 4D, E). Given the four hour difference between the time the *Actβ^ed80^* mutant and the *w^1118^* control reach CW, and considering that *Actβ* mutants grow slower, we calculated that the weight at the time when CW is reached for *Actβ* mutants is similar to that of controls (0.88 mg for *w^1118^* vs 0.84 mg for *Actβ^ed80^*, Fig. 4F). Therefore, we conclude that *Actβ^ed80^* does not affect the CW checkpoint.

Although the CW represents the lower threshold of mass necessary for pupariation without delay, body size can be altered by either a shorter or longer terminal growth period which occurs after CW has been reached (NIJHOUT AND CALLIER 2015). Therefore, we also measured the total time to pupariation. We find that, although *Actβ^ed80^* homozygous mutants grow slowly, they pupariate on average 4 hours earlier when compared to *w^1118^* at 25°C. (Fig. 4G). However, the *Actβ^ed80^/+* heterozygotes also pupariate earlier than *w^1118^* (Fig 4G) suggesting either a haplo-insufficient effect of *Actβ^ed80^* or that this phenotype is attributable to the genetic background differences between *Actβ^ed80^* and *w^1118^*. To examine this issue, we repeated the experiment with two CRISPR alleles. In this case, we find no difference in pupariation timing during the third instar stage between trans-heterozygous *Actβ^4dd^/Actβ^10E^* mutants or *Actβ^4dd^/unc13* heterozygotes compared to *w^1118^* controls (Fig. 4H). However, we also observe that certain allelic combinations such as *w^1118^/unc13* wild-type controls and *Actβ^4dd^/w^1118^* heterozygotes also pupariate 2 hours earlier than *w^1118^* controls (Fig. 4H). Therefore, we conclude that the slight timing differences reflect either differences in genetic background, slight differences in the duration of first and second instar growth, or the limits of timing resolution in this experiment and that *Actβ* loss does not substantially affect developmental timing.

### Overexpression of *Actβ* in its normal pattern produces larger and slower-growing larvae

Since loss of *Actβ* results in small developmentally arrested pupae, we asked whether overexpression of *Actβ* in its endogenous pattern would have the opposite effect on pupal size, viability, and whether it might also affect developmental timing. For this purpose, we overexpressed *Actβ* using an *Actβ-Gal4* promoter enhancer line (ZHU *et al*. 2008; SONG *et al*. 2017a). Relative to either the *UAS-Actβ-3B2* or the *Actβ-Gal4* controls, overexpression of *Actβ* (*Actβ > Actβ-3B2*) results in a significant increase in pupal volume and a delay in developmental timing (Fig. 5A,B). Strikingly, pupariation is delayed over 20 hours compared to either *w^1118^* or *Actβ* mutants however, there is no change in viability (Fig. 5B,C). Together, these data show that *Actβ* regulates body size and perhaps developmental timing in a dose dependent manner.

**Figure 5.**
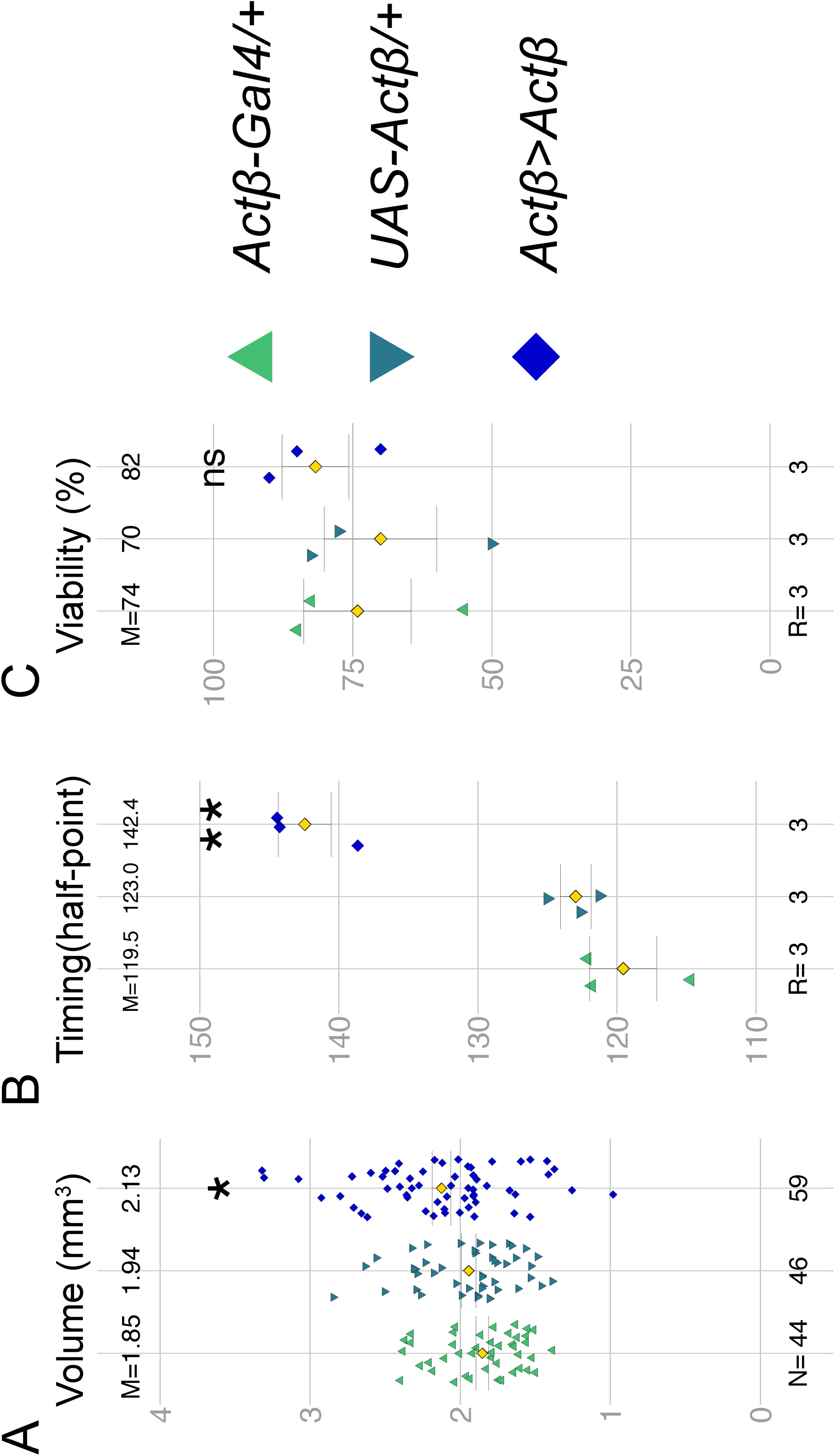
Actβ overexpression increases body size and delays developmental timing. (A) Expression of *UAS-Actβ* using *Actβ-Gal4* significantly increases pupal volume. (B) *Actβ>Actβ* animals pupariate about ∼20 hours later than controls. (C) Adult viability is not significantly impacted in *Actβ>Actβ* animals. M=mean, N=number of individuals, R= number of groups containing 10-30 larva).

### Tissue and cell-type specific rescue experiments identify several potential sources of Actβ for controlling body size

To investigate how Actβ affects body size, developmental timing and viability, we first sought to determine if one or more cell types serves as the source(s) of the ligand that controls different aspects of the mutant phenotype. Several features of endogenous *Actβ* transcription have been previously described including expression in motoneurons, mushroom body neurons, peripheral neurons including multi-dendritic and chordotonal neurons, developing photoreceptors in the eye disc, and in midgut enterocytes (GESUALDI AND HAERRY 2007; TING *et al*. 2007; ZHU *et al*. 2008; KIM AND O’CONNOR 2014; TING *et al*. 2014; MAKHIJANI *et al*. 2017; SONG *et al*. 2017a). We examined the *Actβ* expression pattern in the larvae by crossing an *Actβ-Gal4* to *UAS-cd8GFP* or *UAS-GFP*. We also confirmed expression in particular cell types using RNA *in situ* hybridization (Fig. S3). As previously described, *Actβ* is almost exclusively expressed in the central and peripheral nervous systems (Fig. 6E). More detailed examination reveals that in the central brain lobes, *Actβ-Gal4>UAS-GFP* is expressed strongly in mushroom body neurons and in a 14-cell cluster in the anterior medial region of each brain lobe (Fig. 6A-A’’). A subset of 7 cells within this 14 cell cluster also stain with α-Dilp5 (Fig. 6A’), which marks the ∼7 insulin producing cells (IPCs) (BROGIOLO *et al*. 2001). In the ventral nerve cord, *Actβ* >Gal4 is expressed strongly in the motoneurons, marked by α-p-Mad (Fig. 6G-G’’) (MARQUES *et al*. 2002). We also see strong staining in all α-Dimm-marked neuroendocrine cells (Fig. 6H-H”) (PARK *et al*. 2008).

**Figure 6.**
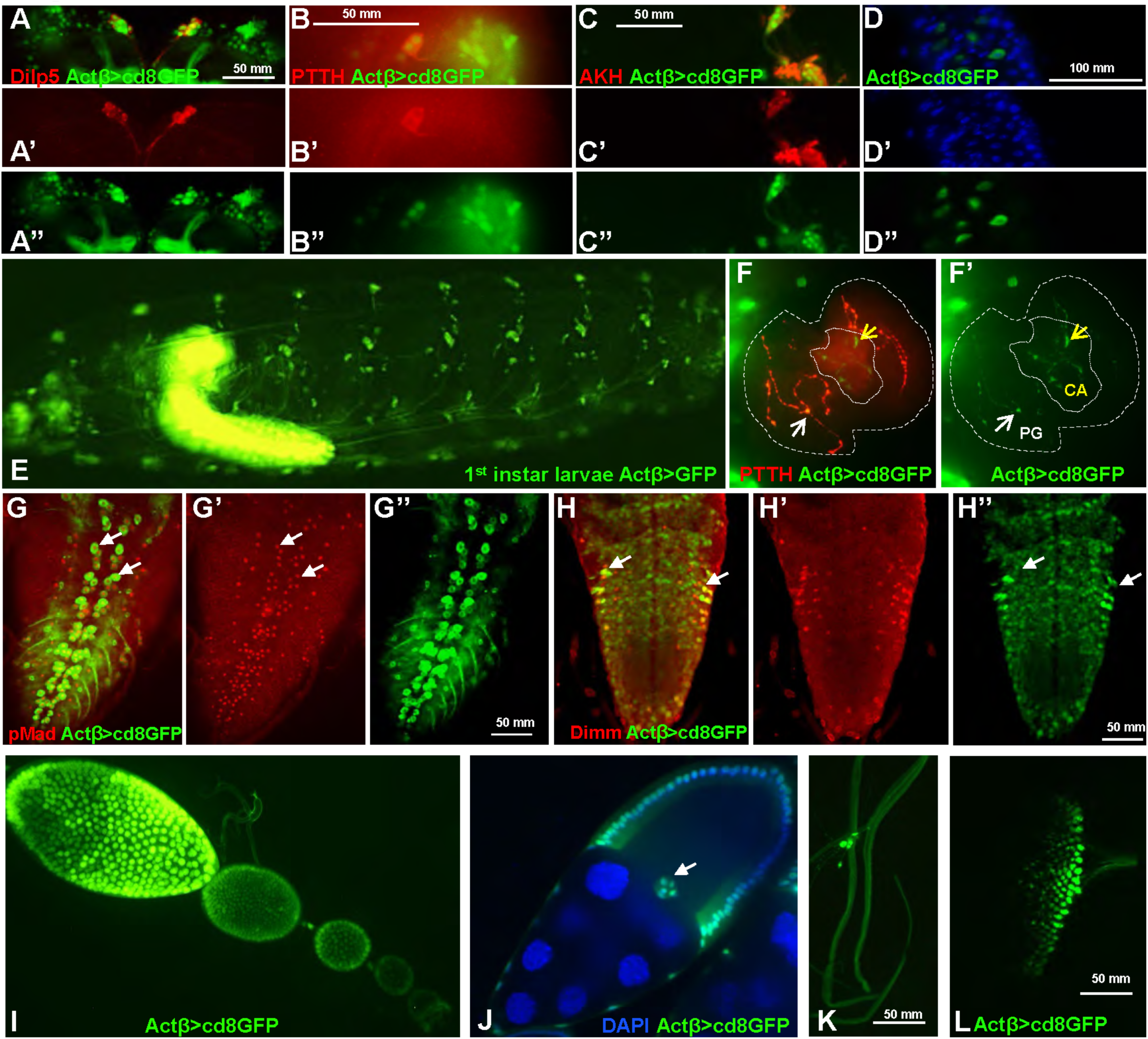
Analysis of *Actβ-GAL4* driver expression pattern (green) in L3 larvae. (A-A’’) Actβ-GAL4 is expressed in the Insulin Producing Cells in the central brain, marked with α-Dilp5 (red). (B-B’’) Actβ reporter is expressed in the cell bodies of PTTH neurons (α-PTTH, red) in the central brain. (C-C’’) Actβ reporter is expressed in AKH producing (α-AKH, red) neurons. (D-D’’) *Actβ-Gal4-*driven GFP is expressed in midgut enteroendocrine cells (Blue DAPI) (E) An intact L1 larvae, *Actβ-Gal4-*driven GFP is expressed in both the central and peripheral nervous systems. (F,F’) *Actβ-Gal4-*driven GFP is found in PTTH synaptic boutons (red) on the PG (thicker dotted line, white arrows) as well as unique boutons in the CA (finer dotted line, yellow arrows). PG= prothoracic gland, CA= corpus allatum. (G-G”) *Actβ* reporter drives expression in the motor neurons (marked with α-pMad red) in the ventral nerve cord (white arrows highlight two individual motor neurons. (H-H”) Actβ reporter is expressed in neuroendocrine cells (α-DIMM, red) in the ventral nerve cord (I) *Actβ-Gal4-*driven GFP is found in follicle cells and the border cells during egg development. (K) *Actβ-Gal4-*driven GFP is found in certain tracheal associated cells and (L) in differentiating photoreceptor cells in the eye disc.

Because of the possible developmental timing defects, we were particularly interested in whether *Actβ* is expressed in the neurons that innervate the ring gland (RG), the major endocrine organ of larvae, or in any of the RG cells themselves. We found strong expression in the corpus cardiacum (CC) cells that produce the hormone Akh which is involved in regulating sugar metabolism (Fig. 6 C-C’’ Fig S3D) (Lee and Park, 2004). While we observed no expression in the cells of the prothoracic gland (PG), which produces the steroid hormone ecdysone (YAMANAKA *et al*. 2013a), or in the corpus allatum (CA) which produces juvenile hormone (RIDDIFORD *et al*. 2010), we did see signal in axons tracts that innervate each of these tissues (Fig 6F-F’). The PG neurons produce prothoracicotropic hormone (PTTH) and innervate the PG portion of the ring gland to regulate ecdysone production (SIEGMUND AND KORGE 2001; MCBRAYER *et al*. 2007). Co-staining of *Actβ>cd8GFP* brains with α-PTTH reveals strong expression in the PG neurons (Fig. 6B-B”). While we have no specific antibody that marks the CA neurons, the GFP positive innervations that we observe on the CA are highly suggestive that the CA neurons express Actβ (Fig. 6F-F’). Actβ is also found in various other unidentified neurons within the central brain and ventral nerve cord. Outside the nervous system, we observe Actβ expression only in a limited number of enterocytes in the midgut (Fig. 6 D-D”) and non neuronal tissue expression includes the enteroendocrine cells (Fig. 6D-D”, Fig S3H) as previously reported (SONG *et al*. 2017a), and the ovariole follicle and border cells (Fig. 6 I and J, Fig. S3G) and some tracheal cells. (Fig. 6K, Fig. S3F). Our observation that the rare escaper females are fertile suggests that Activin signaling in the follicle cells is either not required for full fertility or that its expression might be redundant with another activin like ligand such as Dawdle or Myoglianin.

### Motoneuron-derived Actβ regulates body size and viability

To determine which *Actβ*-expressing cell types influence size and viability, we attempted rescue experiments using different tissue specific Gal4 drivers to overexpress the *Actβ* transgene in the *Actβ^ed80^* mutant background. *Actβ^ed80^* mutants with one copy of either the *UAS-Actβ* or the various *Gal4* transgenes served as negative controls. Since overexpression of *Actβ-GAL4* driving *UAS-Actβ* is sufficient to increase body size (Fig. 5A), here we asked whether it is able to rescue the small body size (pupal volume) and pupal lethality of *Actβ* mutants. Indeed, *Actβ>Actβ* in the mutant background is not only sufficient to rescue body size, but actually produces larger animals (47% bigger Fig. 7A) similar to what we see upon overexpression in a wild type background (Fig.5A). Overexpression of Actβ in its normal pattern also resulted in strong rescue of lethality (Fig. 7B, 42.7% viability vs 1-4% viability of mutant controls; the test cross viability rate for *w^1118^* is 73.8%, Fig. 7B).

**Figure 7.**
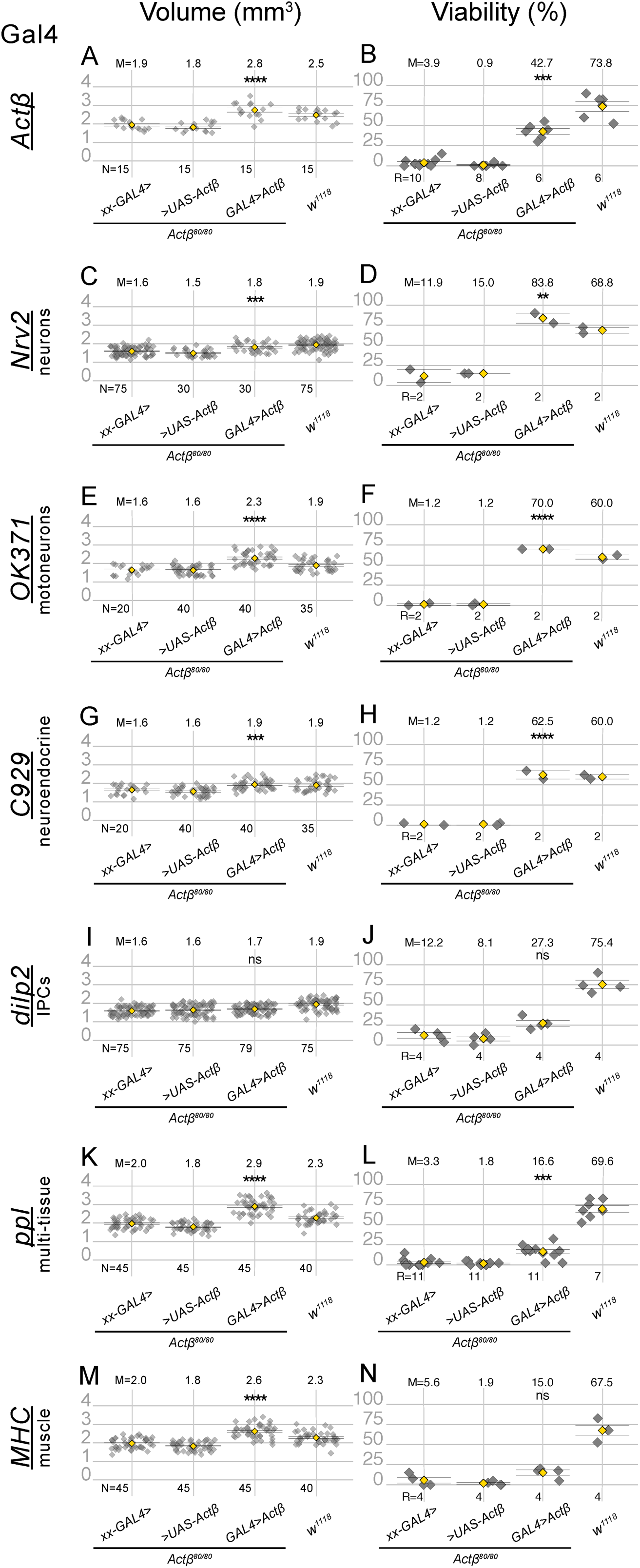
Expression of Actβ from specific cell types differentially rescues Actβ mutant phenotypes. The first two groups in each panel are controls in which *Actβ* mutants contain one copy of either the GAL4 driver (indicated on the left side of each panel row) or the UAS Actβ transgene. The third group in each panel is the testcross, and the last group in each panel is the *w^1118^* control. All GAL4 drivers (except *dilp2-GAL4)* used to overexpress *Actβ* rescue body size phenotype (A, C, E, G, K, M). Overexpressing *Actβ* in neuronal tissues (D, F, H) completely rescues adult viability phenotype, and partially rescue when overexpressed in body wall muscles using MHC-GAL4 (L). ANOVA was used to determine statistical significances between genotypes with one copy of either Gal4 or UAS transgenes in an *Actβ^ed80^* homozygous background compared to animals with both Gal4 and UAS transgenes in an *Actβ^ed80^* homozygous background. *w^1118^* is the wild-type control and was reared side-by-side in each case. M=mean, N=number animals, R= repetition number (10-30 animals per rep).

To narrow down the relevant source of ligand that regulates each phenotype, we used increasingly restrictive (tissue specific) Gal4 lines to overexpress *Actβ* and then measured body size and viability. Overexpression of *Actβ* using the pan-neuronal driver, *nrv2-Gal4*, rescues both body size and adult viability (Fig. 7C and D). Surprisingly, overexpression of *Actβ* from the motoneurons, *OK371>GAL4,* alone rescues both body size and viability (Fig. 7E and F). Interestingly, *Actβ* overexpression in Dimm^+^ neuroendocrine cells (*C929>Gal4*), also rescues both body size and viability (Fig. 7G and H). Just like overexpression of *Actβ* from its endogenous sources, we also found that overexpression of *Actβ* in either motoneurons or neuroendocrine cells in wildtype animals also produces large adults (Fig. S4). Lastly, overexpression of *Actβ* in only the IPCs (*dilp2>Gal4*), which makes up a much smaller subset of all neuroendocrine cells, does not rescue either phenotype (Fig. 7I and J).

The finding that expression in only the motoneurons rescues body size suggests that Actβ may be supplied directly to the muscles via the neuromuscular junctions. However, we also find that overexpression in neuroendocrine cells is sufficient to rescue body size, which suggests that Actβ may be able to function as a systemic endocrine signal and need not be directly delivered to the muscle via the neuromuscular junction synapse. Therefore, we asked if expression of *Actβ* from non-neuronal, but highly secretory tissues, was able to rescue various aspects of the null phenotype. Interestingly, expression of *Actβ* using the *ppl-Gal4* (fat body and muscle) driver increases pupal volume beyond wild-type levels and partially rescues adult viability (Fig. 7K and L). Overexpression in only the body wall muscles (*MHC-Gal4*) also increases body size beyond wild type levels, but does not rescue adult viability (Fig. 7M-N). However, we note that overexpression of *Actβ* using either *MHC-Gal4* or *ppl-Gal4* in a wild type background results in most animals dying as large oversized and curved pupae (Fig. S5). These phenotypes are likely due to hyperactivation of TGFβ signaling in the muscles because we observe a similar phenotype when a constitutively activated version of Babo is overexpressed in the muscles (Fig. S5). Taken together, these results suggest that, although Actβ signaling in muscles is required for proper body size, too much signaling in muscles can be deleterious. We were not able to specifically test the ability of enteroendocrine-derived Actβ to rescue mutant phenotypes, because overexpression of *Actβ* using the midgut enteroendocrine cell driver (*EE-Gal4*) (SONG *et al*. 2017a) is lethal in both wild type and *Actβ* mutant backgrounds, likely due to overexpression in many cells besides enteroendocrine cells, including fat body, CNS and PNS (data not shown). In summary, we conclude that since overexpression of Actβ from motoneurons or neuroendocrine cells rescues both body size and viability and can increase body size when overexpressed from these sources in wild type animals, they are likely the most important endogenous sources of ligand for viability and body size control.

### Motoneuron derived Actβ signals through the canonical Babo/dSmad2 pathway to control muscle and body size

The rescue experiments described above suggest that either motoneurons or Dimm^+^ neuroendocrine cells, or both can produce enough Actβ to regulate body size. Since data from overexpression alone does not reflect the *in vivo* importance of various endogenous ligand sources, we sought a complementary set of loss-of-function data using tissue specific RNAi knockdown. First, we tested all publicly available (TRiP, VDRC, and NIG) *Actβ* RNAi lines to phenocopy the *Actβ* mutant. Using the ubiquitous driver *da-GAL4* to overexpress *dicer2* along with the various *Actβ* RNAi lines, we find that only the TRiP stock (BDSC#29597) can phenocopy the small, dead pharates similar to *Actβ* null alleles (data not shown). Most other lines produce viable flies of normal size suggesting that they are not very effective in knocking down endogenous *Actβ.* Both NIG lines (1162R-1 and 1162R-2) produce a more severe phenotype (early larval lethality) compared to the null suggesting it may have off-target effects.

Using the TRiP 29597 RNAi line, we tested whether knockdown of *Actβ* in either all neurons, motoneurons or neuroendocrine cells alone phenocopies any aspect of null alleles. We find that knockdown in Dimm**^+^** neuroendocrine cells (*C929-Gal4*) produces viable normal-sized flies (data not shown). In contrast, knockdown in all neurons (Elav>Gal4 Fig. 8A) or motoneurons (*OK371-Gal4*, Fig. 8A,B) completely phenocopies *Actβ* nulls giving rise to small, dead pharates with rare escapers that hold out their wings and have a slow gait (Movies 3,4). The *OK371* driver is not expressed in the Dimm**^+^** neuroendocrine cells (Fig. S6) leading us to conclude that the motoneurons are the major source of endogenous *Actβ* that regulates body size and viability.

**Figure 8.**
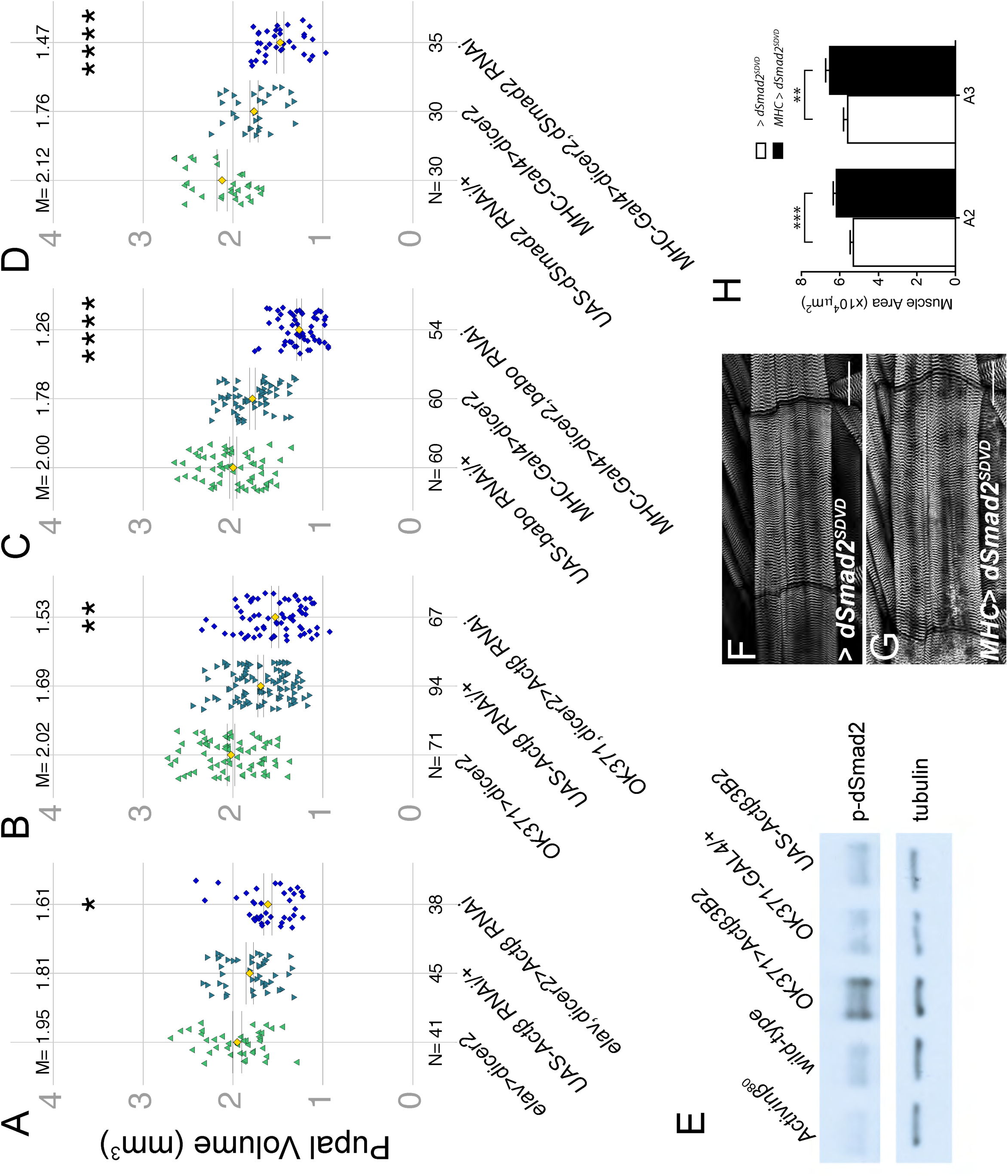
Motor neuron derived Actβ signaling through Babo and dSmad2 controls body size. (A). Knockdown of *Actβ* using a pan-neuronal driver (*elav-Gal4*) with *UAS-dicer2* reduces pupal volume (B) Motor neuron knockdown of *Actβ* using *OK371-Gal4* to also reduces pupal volume. (C) Knocking down the Actβ receptor *baboon* or the signal transducer *dSmad2* (D) in muscles using *MHC-GAL4* with UAS-*dicer2* reduces pupal volume. M=mean, N=number (E) *Actβ* mutants have lower levels of p-dSmad2 (E, lane 1 vs 2) in carcass tissue lysates (cuticle, skeletal muscle). While overexpressing Actβ in motor neurons leads to increased level of dSmad2 compared to controls (E, lanes 3-5) Anti-tubulin staining serves as a loading control. (F – H) Overexpressing activated dSmad2 (dSmad2^CA^) in the muscles using *MHC-GAL4* increases muscle size by ∼20%.

We next determined if motoneuron-derived *Actβ* signals to the muscle via the canonical Smad2 pathway. Canonical TGFβ signaling is mediated by the Activin receptor, Babo, and the signal transducer, dSmad2. In muscles, overexpressed Babo is localized to the postsynaptic neuromuscular junction, perhaps sensitizing the muscles to receive motoneuron derived Actβ (KIM AND O’CONNOR 2014). Indeed, RNAi knockdown of *Babo* or *dSmad2* in the body wall muscle results in smaller pupal volume (Fig. 8C and D). Furthermore, on Western blots, we detect lower levels of phosphorylated dSmad2 in *Actβ* mutant carcass extracts (containing somatic muscle, cuticle and associated cells) compared to the *w^1118^* control (Fig. 8E). Conversely, overexpression of *Actβ* in motoneurons both increases pupal/adult size (Fig. S4) and p-Smad2 levels in the carcass (Fig. 8D-E), which is similar to overexpression of activated dSmad2 in the muscle, producing large larval muscles (Fig. 8F-H) or overexpression of activated Babo or dSmad2 results in adult flies with extended abdomens (Fig. S7A-C). Interestingly, the *MHC>dSmad2(SDVD)* animals with larger body size have slightly smaller wings, not larger wings as expected if organs are actively scaled to maintain size proportion between muscles and appendages (Fig. S7D).

## Discussion

Identifying and characterizing how inter-organ signals regulate physiologic and metabolic homeostasis during development and adulthood is of central importance. Various types of inter-organ signals are also likely to be necessary for coordinating growth between organs during development to achieve proper body proportions (DROUJININE AND PERRIMON 2016). In this report, we demonstrate that Actβ is a key brain-derived factor that regulates somatic muscle size in *Drosophila* by signaling through the canonical Smad-dependent pathway. Furthermore, we find that disruption of Actβ signaling alters larval and adult organ allometry suggesting that Actβ might be a component of an inter-organ signaling pathway that helps coordinate muscle growth with appendage growth.

### Localized versus systemic effects of Actβ

The question of whether Actβ acts locally or systemically via the hemolymph to target tissues is an important issue raised by our study and previous work (SONG *et al*. 2017a; SONG *et al*. 2017b). On the one hand, we find that *Actβ* is strongly expressed in most if not all neuroendocrine cells. We also find that overexpression of *Actβ* from these cells results in the rescue of mutant phenotypes and overgrowth of wild type animals indicating that direct tissue contact is not necessary for Actβ signaling to control muscle size. However, we also find that depleting *Actβ* expression in just the motoneurons phenocopies *Actβ* mutants while depletion in neuroendocrine cells does not do so, at least with the *929>Gal4* driver. Therefore, we conclude that, while high systemic concentrations of Actβ produced by overexpression is capable of regulating muscle growth, the endogenous systemic level supplied by the combination of the neuroendocrine and the enteroendocrine cells is not sufficient to do so.

Whether local or systemic Actβ signaling is important in other contexts is less clear. Interestingly, Actβ has also been implicated in regulating hemocyte proliferation and adhesion within hematopoietic pockets localized on the larval body surrounding a number of peripheral neurons that express *Actβ* including da and chordotonal peripheral neurons (MAKHIJANI *et al*. 2017). In contrast, enteroendocrine derived Actβ is able to affect AKH receptor levels in the fatbody to regulate glycemic index on a high sugar diet (SONG *et al*. 2017a). In addition, it has been reported that upon mitochondrial perturbation, muscle-derived Actβ signals to the fat body to regulate triglyceride levels (SONG *et al*. 2017b). In either case these observations raise the question: what dictates the requirement of a local versus a systemic signal for Actβ function? In the muscle motoneuron synapses and hematopoietic pocket paradigms, there may be physical barriers that help concentrate ligand from a local source to levels sufficient to produce a response. In the case of muscles, motoneuron synapses are embedded within the muscle fiber (PROKOP 2006; PROKOP AND MEINERTZHAGEN 2006) and therefore delivery of Actβ directly to the NMJ via the synapse likely provides a highly effective signal, especially since its receptor Babo is also highly concentrated at the postsynaptic NMJ (KIM AND O’CONNOR 2014). This possibility might also account for the discrepancy between our findings (Fig. S3) that muscles do not express Actβ under normal conditions, while mitochondrial perturbations in muscle appears to release Actβ for signaling to the fat body (SONG *et al*. 2017b). Perhaps disturbing mitochondrial function in muscle disrupt synaptic structure such that Actβ is liberated from defective NMJ synapses. Likewise, the hematopoietic pockets might provide a similar restricted niche-like signaling environment that is able to modulate hemocyte proliferation and adhesion. These types of physical constraints may limit the ability of endogenous circulating Actβ to produce sufficient levels of signaling at these locations, except under overexpressed conditions.

Another factor influencing the cellular response to Actβ levels is the composition of the surface receptors. The *babo* locus produces three receptor isoforms that only differ in the extracellular ligand binding domain, and each likely has a different affinity for the three Activin-like ligands (JENSEN *et al*. 2009; UPADHYAY *et al*. 2017). Therefore, the complement of receptor isoforms on a cell’s surface is apt to determine the sensitivity of the cell or tissue to Actβ signals.

### Mechanisms of Actβ control of tissue size

The molecular mechanism(s) by which Actβ affects tissue growth are unclear. The most well characterized factors that regulate insect body size are all systemic signals such as juvenile hormone, ecdysone and the IIS/TOR pathways (REWITZ *et al*. 2013; MIRTH AND SHINGLETON 2014; BOULAN *et al*. 2015; KOYAMA AND MIRTH 2018). For example, *Ptth* mutants delay ecdysone accumulation allowing larvae to grow for an additional 24 hours ultimately leading to larger flies (SHIMELL *et al*. 2018). In this report, we demonstrate that Actβ, although it is expressed in the PTTH-producing neurons, does not appear to affect ecdysone signaling since *Actβ* loss affects neither critical weight nor developmental timing.

Interestingly, we also see Actβ positive innervation of the CA organ which produces juvenile hormone (JH), and lowering JH levels in Drosophila leads to production of smaller flies by slowing the overall growth rate (RIDDIFORD *et al*. 2010.; MIRTH *et al*. 2014) Since we also observe a slower growth rate in *Actβ* mutants it is possible that Actβ might work to slow growth via reduction of JH signaling. However, given the strong expression of Actβ in all Dimm^+^ neuroendocrine cells which secrete numerous peptides that regulate many aspects of behavior and metabolism including insulin, many other mechanisms must be considered for slowing growth. While food intake does not seem to be altered in Actβ mutants, it is possible that nutrient absorption or metabolic flux is disrupted. The latter possibility is particularly attractive since we see strong expression of *Actβ* in the CC organ, which produces the Drosophila glucagon-like hormone (Akh), and in the insulin producing cells (IPCs) in the brain. As previously noted, Actβ has been implicated in regulating AkhR levels in the fat body (SONG *et al*. 2017a) and perhaps it may also influence Akh synthesis or release. Furthermore, Dawdle, another Drosophila Activin-like ligand that also signals through dSmad2 has been previously shown to regulate metabolism and carbohydrate utilization (CHNG *et al*. 2014; GHOSH AND O’CONNOR 2014). Therefore, Actβ signaling through dSamd2 may also regulate global carbohydrate synthesis or aspects of metabolism to adjust the larval growth rate.

Regardless of how overall growth defects occur, it is important to remember that not all larval tissues respond equally to Actβ. The brain and imaginal discs, for example, are of normal size while fat body and muscle are significantly smaller. In addition, the size and viability defects can be largely rescued by expression of Actβ solely in motoneurons. Therefore, it seems unlikely that a primary defect in systemic levels of insulin or Akh would account for the tissue specific responses. Rather, it is likely that alterations in muscle metabolism and perhaps factors secreted by muscles could account for the small muscle/body size.

### Larval versus Adult requirements for Actβ

The requirement of Actβ for adult eclosion raises several issues. The first is whether the low eclosion rate is primarily a muscle defect or a neuronal problem since both must be coordinated to produce the complex set of motor behaviors required for eclosion. Interestingly, ablation of Dimm+, Eclosion hormone (EH) producing neuroendocrine cells, (PARK *et al*. 2008), results in a defective eclosion motor program, which involves a series of coordinated head, thorax, and abdominal muscle contractions that ejects the animal through the operculum and out of the pupal case (MCNABB *et al*. 1997). It may be that the small adult muscles lack the power to properly execute the eclosion motor program. In addition, the small muscle phenotype may also partially explain why the *Actβ* mutant adult escapers walk slowly and cannot move their wings. However, this must be reconciled with the observation that *Actβ* mutant larvae exhibit no obvious defect in locomotion, even though they have a similar proportional reduction in overall body and muscle size.

Improper synaptic development or NMJ function could also potentially account for adult locomotion defects. However, we have previously shown that, at least in larvae, the NMJ size and bouton number are not affected in *babo* and *dSmad2* mutant larvae when normalized to the smaller muscle size (KIM AND O’CONNOR 2014). Nevertheless, we did uncover a number of electrophysiological alterations including a decrease in the number and frequency of miniature excitatory potentials and a depolarized muscle membrane resting potential, both of which were primarily attributed to defective Actβ signaling in muscles (KIM AND O’CONNOR 2014). Despite these defects, the large action potentials in *babo* and *dSmad2* mutants are relatively normal and, as described, there are no obvious larval locomotion defects (KIM AND O’CONNOR 2014). Since adult muscles are formed *de novo* during metamorphosis, it is possible that during this time more extreme defects in muscle or neuron physiology develop in *Actβ* mutants, perhaps leading to a more strongly depolarized muscle, for example, that would interfere with proper muscle function.

The motoneuron source of Actβ also raises questions concerning whether Actβ production/release is muscle/neuron activity dependent. We find that overexpression of Actβ in motoneurons can produce bigger muscles, but whether increased muscle activity also accompanies higher Actβ expression/secretion triggering increased muscle growth is an interesting issue to address. We note, however, that adult muscles, which develop during the immobile pupal stage, are also likely smaller than wild type in *Actβ* mutants suggesting that significant muscle activity is not likely required for Actβ release.

### Body-appendage scaling

One of the more novel features of the *Actβ* null phenotype is the disproportionate effect it has on muscle size compared to other tissues. One might expect that evolutionary pressures fine tune mechanisms to coordinate muscle size with the size of the appendage that it moves. This is perhaps especially true in winged insects where flight muscle and wing size should be coordinated to produce efficient flight. Such coordination between wing and body size in response to environmental perturbations has been best studied in *Manduca sexta* (NIJHOUT AND GRUNERT 2010; NIJHOUT AND CALLIER 2015). In this insect, nutritional restriction can result in as much as a 50% reduction in body size, with the wing scaling proportionally and containing half as many cells (NIJHOUT AND GRUNERT 2010). This scaling mechanism utilizes a shift in the amplitude and kinetics of steroid hormone production during the last instar stage. Since this mechanism involves systemic factors that adjust the growth rate of the whole body, presumably affecting muscles and discs simultaneously, it does not really address whether specifically perturbing muscle growth can directly or indirectly affect growth of the wing or other appendages.

In Drosophila, alteration in the growth properties of one imaginal disc perturbs growth of other wild type discs in a coordinated manner so that adults emerge with properly proportioned structures (SIMPSON AND SCHNEIDERMAN 1975; SIMPSON *et al*. 1980; STIEPER *et al*. 2008). Once again, the inter-organ signaling mechanism involves alteration in the levels of systemic hormones (PARKER AND SHINGLETON 2011; MIRTH AND SHINGLETON 2012; GOKHALE *et al*. 2016). In these reports, is not clear whether muscle size was also altered to produce isometric scaling between it and the imaginal discs. However, it is interesting to note that growing Drosophila at low temperatures produces hyperallometric scaling where the wing size is disproportionally larger relative to body size (SHINGLETON *et al*. 2009). Since we observe similar phenotypes in *Actβ* mutants grown at normal temperatures, it is intriguing to speculate that Actβ signaling might mediate hyperallometric scaling between wing and body in response to temperature.

The only other report that we are aware of where Drosophila larval muscle size was specifically manipulated, and the effect on growth of other tissues examined, involved genetic alteration of insulin signaling (DEMONTIS AND PERRIMON 2009). Similar to our analysis of *Actβ*, the level of insulin signaling in muscle is directly correlated with muscle, appendage, and overall animal size. Nevertheless, our findings for Actβ show several notable differences. First, insulin gain-of-function signaling in muscle lead to larger bodies and larger wings (DEMONTIS AND PERRIMON 2009), while we find that increased Actβ signaling in muscles results in larger bodies, but slightly smaller wings. In the insulin loss-of-function case, both muscles and wings were smaller, the latter due to a reduction in cell size not cell number. However, in the case of *Actβ* mutants, we see only a 4% decrease in wing size with no change in cell size. In both cases, the effect on muscle size appears to be much more dramatic than the effect on appendage size. Therefore, if a scaling mechanism exists, then either insulin or Actβ loss disrupts it, or it is not isometric as is found for the nutrient-dependent body-wing scaling response in *M. sexta*. The general similarity in phenotypes produced by insulin or Actβ signaling suggests that Actβ may exert its effect on muscle and body size, in part, through the insulin signaling pathway, a possibility that we are currently exploring.

### TGFβ control of body size in other animals

TGFβ regulation of body or muscle size has been reported in both *C. elegans* and mammals. In *C. elegans*, BMP-type factors are also secreted from a specific set of neurons and appear to act systemically to regulate the size of the hypodermis through a canonical Smad dependent pathway (TUCK 2014). In fact, the term Smad is a compound word derived from the *C. elegans* gene *sma* meaning small, and the Drosophila gene *mad* (<u>m</u>others <u>a</u>gainst <u>d</u>pp) which were the founding members of the Smad family of TGFβ signal transducers (DERYNCK *et al*. 1996). Recently, many transcriptional targets for the Sma factors in *C. elgans* have been identified, among which are several collagens that are major structural components of the hypodermal body-wall (MADAAN *et al*. 2018). Hence, we speculate that one set of targets for both Actβ and insulin signaling in the *Drosophila* muscle could be structural proteins that build muscle. It is also possible that target gene expression is indirectly regulated by DNA copy number. In Drosophila, larval and adult muscle are polyploid tissues where DNA content is controlled by endocycling. Both Actβ (this report) and insulin signaling (DEMONTIS AND PERRIMON 2009) appear to regulate nuclear size in many polypoid tissues, possibly indicating that control of the endocycle maybe the primary mechanism regulating tissue size. However, at least in Actβ mutants not all polyploid tissues show regulation in the same direction, i.e. muscle, fatbody and the prothoracic gland all show smaller nuclei while the salivary gland has larger nuclei. Whether systemic Actβ signaling is directly regulating size of polyploid tissues or acts indirectly through a muscle-derived myokine needs to be determined.

In mammals, the best characterized example of a TGFβ-type factor that regulates body and muscle size is provided by Myostatin (Mstn). Loss of Mstn was discovered to cause the muscle overgrowth phenotype of Belgian blue cattle and subsequent work in many other species including humans confirmed that Mstn is a negative regulator of muscle mass (MCPHERRON *et al*. 1997; MCPHERRON AND LEE 1997). *Mstn* mutant muscles have both an increase in myofiber number (TRENDELENBURG *et al*. 2009; MATSAKAS *et al*. 2010) as well as myofiber size (MCPHERRON AND LEE 1997; ELASHRY *et al*. 2009). The molecular basis for the phenotype appears to be an alteration in protein homeostasis, where proteasome and autophagic degradative capacity is reduced relative to protein synthesis (LEE *et al*. 2011; LOKIREDDY *et al*. 2012). Mstn signals through Smads2/3 and is therefore considered to be within the TGFβ/Activin subgroup in the TGFβ superfamily. Additional studies of Activin ligands themselves suggest that that they also act as negative regulators of muscle mass similar to Mstn (ZHOU *et al*. 2010). Furthermore, studies on the role of BMP signals in muscle size control suggest that they function as dominant positive regulators of muscle mass by promoting protein synthesis instead of breakdown (SARTORI *et al*. 2013; WINBANKS *et al*. 2013).

Our present work shows that in Drosophila, Actβ is a positive regulator of muscle mass, by affecting myofiber size not number. It is worth noting that Drosophila has a close Myostatin homolog that is called *myoglianin* (*myo*) and recent studies suggest that loss of *myo* in muscle produces larger fibers similar to the vertebrate homolog (AUGUSTIN *et al*. 2017). How Actβ and Myo interact will be interesting to examine as will the role for BMPs in *Drosophila* muscle size determination. Lastly, the question of whether Mstn loss in vertebrates affects scaling of other tissues is largely unexplored although it does appear that bone density is increased in *myostatin* mutant animals (ELKASRAWY AND HAMRICK 2010). Additional studies of how local versus systemic roles of TGFβ ligands might affect growth and scaling between tissues and organs in vertebrates should be enlightening.

## Materials and Methods

### Fly lines

For overexpression experiments single copies of Gal4 and UAS transgenes were used. *Actβ-Gal4* and *UAS-Actβ* (3B2) were previously described (ZHU *et al*. 2008). *C929-Gal4*, dilp2*-Gal4*, *Elav-Gal4*, *Mef2-Gal4*, *MHC-Gal4*, *Nrv2-Gal4*, *OK371-Gal4*, *ppl-Gal4*, *UAS-dicer2, UAS-cd8::GFP,* and *UAS-Actβ RNAi Ok6>Gal4* were all from the Bloomington stock center. *UAS-babo RNAi*, *UAS-dSmad2 RNAi*, were from O’Connor lab stocks. (details of construction available upon request). *UAS-dSmad2^SDVD^* (constitutively activated dSmad2) was previously described (GESUALDI AND HAERRY 2007).

The *Actβ*^ed80^ allele is an EMS-induced substitution leading to a premature stop codon and presumed to be a null mutation (ZHU *et al*. 2008). The chromosome carrying the *Actβ*^ed80^ allele (fourth) also contains a variegating *w*^+^ transgene (P{hsp26-pt-T}39C-12, FlyBaseID= FBti0016154) inserted between *Hcf* and *PMCA* (John Locke, personal communication). This *w^+^*transgene causes red speckles with dominant inheritance in an otherwise *w^-^* background.

*Actβ^4E^* and *Actβ^10E^* and *Actβ^4dd^* were all generated using the CRISPR/Cas9 system. Two guide RNAs were cloned into the BbsI site of pU6-BbsI-chiRNA plasmid (obtained from Addgene) and injected by Best Gene into *w^1118^; PBac{y*[*+mDint2*]*=vas-Cas9}VK00027* on chromosome 3 (Bloomington Stock Center #51324). The following guides were used to target the genomic locus, guide 1: 5’-GGGTTGTGGAAATGACTTCC-3’, guide 2: 5’-GCGATTGCACGGGCTCTTTT-3’. G0 male flies were backcrossed to a balancer stock (*CiD/unc13-GFP*) to isolate *w^1118^;;;Actβ^?^/unc-13-GFP* stocks. To identify new *Actβ* alleles, DNA from homozygous (non-GFP) larvae was used to PCR amplify the genomic region flanking the CRISPR target sites using the following primers (FWD: 5-CTGCTGCAACAGCCTTGGCTCCC-3; REV: 5-GGGGCGCAACACGGTCGCATTCC-3). Line 4E and 4dd are independent ∼3 kb deletions that remove that remove exons 2 and 3. Line 10E is a ∼1.3 kb deletion that removes exon 4 and 5. Exact deletion junction sites are available upon request.

### Rearing Conditions

Eggs were collected over a 2-3 hour time period on apple juice plates inoculated with yeast paste and aged until hatching into first instar larvae. Larvae of the desired type were then transferred to vials containing standard cornmeal food (Bloomington recipe) and incubated at 25°C in a 12 hour light/dark cycle until scoring. Animals were transferred to vials at a low density (30 or 40 per vial) to prevent crowding affects.

### Size measurements of larval tissues and nuclei

To measure size of larval organs, tissues were prepared using standard protocols for immunohistochemistry (see below). To measure size of larval body wall muscles, larval fillets of late wandering L3 larvae were prepared, and the surface area of muscle #6 of the A2 segment was measured in FIJI by outlining the muscle segment using the free-hand selection tool. Larval brains were stained with DAPI and rhodamine-phalloidin and placed onto a glass microscope slide between two #2 coverslips that acted as a bridge to prevent deforming the shape of the brain lobes. Confocal Z-stacks of the entire lobe were captured, and manual 3D segmentation using ITK-SNAP (PMID: 16545965) was used to measure lobe volume. Imaginal discs were stained with DAPI, imaged using confocal microscopy, and then maximum intensity projections were generated and processed in FIJI using the threshold and measure functions to obtain a 2D area of each disc. For fat body, proventriculus, muscle salivary and the prothoracic glands, tissue was stained with DAPI and rhodamine-phalloidin and then Z stacks obtained. Nuclear size was measured using FIJI (SCHINDELIN *et al*. 2012) at the sections where nuclei were largest.

### Pupal Volume determination

Pupal volume was calculated from the length and width of individual pupae assuming a prolate spheroid shape [V = (4/3) π (width/2)2 (length/2)] (DEMONTIS AND PERRIMON 2009). Pupal length was measured from the anterior tip midway between spiracles to the base of the posterior spiracles. Pupal width was measured at the mid-length of the pupae.

### Measurement of Adult appendage sizes

Adult specimens were fixed in 95% Ethanol. Structures were dissected and mounted in Canadian Balsam (Sigma, C1795) and Wintergreen oil (Sigma, M2047) solution (50:50). To measure size (length or area) of adult body parts images were processed in FIJI using either the free-hand or polygon tool (illustrated by red lines in Fig. 2).

### Developmental Timing and Growth Assay

To measure developmental timing, flies were transferred to a constant light environment for at least 2 days prior to egg lay and all subsequent assays were carried out under constant light conditions to avoid circadian rhythms. Eggs were collected on apple juice plates with yeast paste for two hours. The next day, early L1 larvae were transferred to standard cornmeal food with yeast paste. For *Actβ^ed80^/Actβ^ed80^* mutants the time to midpoint of pupariation was measured. For the CRISPR alleles, an additional synchronization step was employed. After ∼48 hours of growth, 20-30 synchronized L2-L3 ecdysing larvae were transferred to cornmeal food without yeast paste to measure time to pupariation. Pupariation was scored every 2 hours by monitoring for anterior spiracles eversion and larval movement. The half point is the time it takes for half of the population to pupariate, which is calculated using a simple linear regression.

To measure growth rate, L3 larvae were cultured for appropriate times after L2-L3 ecdysis, washed in water and weighed individually on a Mettler Toledo XP26 microbalance. For adult mass, groups of 8-10 animals were weighed on the microbalance.

### Statistics

Data were analyzed using either Graph Pad – Prism or R-studio. A single test variable was compared to a single control using a Welch Two Sample t-test. Multiple test variables were compared to controls using a one way ANOVA followed by Tukey’s multiple comparison test. For rescue experiments with two controls and one test cross, the test cross must be significantly different in the same direction (e.g. larger) to be considered a significant result. Where the test cross was reported to be *x* units different than the controls, the different was in reference to the control with smaller variation. P values designation: ns= not significant, * p<0.05, ** p<0.01, *** p<0.001, **** p<0.0001.

### Immunohistochemistry

Wandering third instar larvae were rinsed, dissected, fixed in 3.7% formaldehyde in PBS for 25 minutes, and then washed three times in PBS-(0.1%) TritonX-100. Samples were incubated with primary antibody overnight at 4°C followed by secondary antibodies for 2 hours at 25°C. Tissues were mounted in 80% Glycerol. The following stains and antibodies were used: Rhodamine-phalloidin (Molecular Probes R415), α-Dachshund (DSHB, mAbdac2-3), α-PTTH (guinea pig) a gift from P. Leopold, α-p-Mad (Eptitomics), α-DIMM a gift from P. Taggert.

### Microscopy

Confocal images were generated using a Zeiss Axiovert microscope with a CARV attachment or Zeiss LSM710. Pupae, adult heads, and body were imaged live with Zeiss Stemi stereo microscope using a 1X objective. Adult wings and legs were imaged using Nikon Optiphot light microscope with a 4X objective. Trichomes were imaged using a 40X objective.

### Western Blots

L3 larvae were dissected and all organs were removed from the carcass samples. Carcass samples were lysed with reducing gel loading buffer. Bands were resolved on 4-12% gradient gels (Invitrogen), and transferred to PVDF membrane (Biorad). Membrane blocking and antibody incubation were performed using standard protocols for ECL detection. α-pSmad2 (CST, 138D4) and α-tubulin (Sigma, T9026) were used at 1/1000 dilutions. Bands were visualized using Pierce ECL Western Blotting Substrate (#32209).

### Data Availability Statement

Strains and plasmids are available upon request. Supplemental files available at FigShare. Movie S1 illustrates the defective shock response of Actβ escaper females compared to heterozygous controls. Movie S2 shows a close up view of Actβ mutant females exhibiting poor locomotion and a held out wing phenotype compared to heterozygous controls. Movie S3 demonstrates a defective shock response of adults in which Actβ was knocked down in motoneurons using RNAi (Ok371>Gal4, UAS Actβ RNAi). Movie S4 shows a close up view of Ok371>Gal4, UAS Actβ RNAi Actβ knockdown adults exhibiting poor locomotion and a held out wing phenotype similar to that exhibited by Actβ null escaper flies (Movie 1).

## Acknowledgements

We would like to thank: Aidan Peterson, MaryJane O’Connor, and Heidi Bretscher for comments on the manuscript; David Zhitomirsky for making guide RNA constructs to generate CRISPR/Cas9 alleles, P. Leopold for the anti-PTTH, P. Taggert for the anti-Dimm antibody, M. Titus for providing the microscope for live imaging of feeding larvae, and the Bloomington Drosophila Stock Center for providing numerous fly lines.

## Author Contribution

Conceptualization: M.B.O., L.M.T., experimentation and data analysis: L.M.T., A.U., X.P., M.J.K., M.B.O., writing: M.B.O., A.U., L.M.T., funding acquisition: L.M.T., M.B.O.

## SUPPLIMENTAL FIGURE LEGENDS

**Figure S1.**
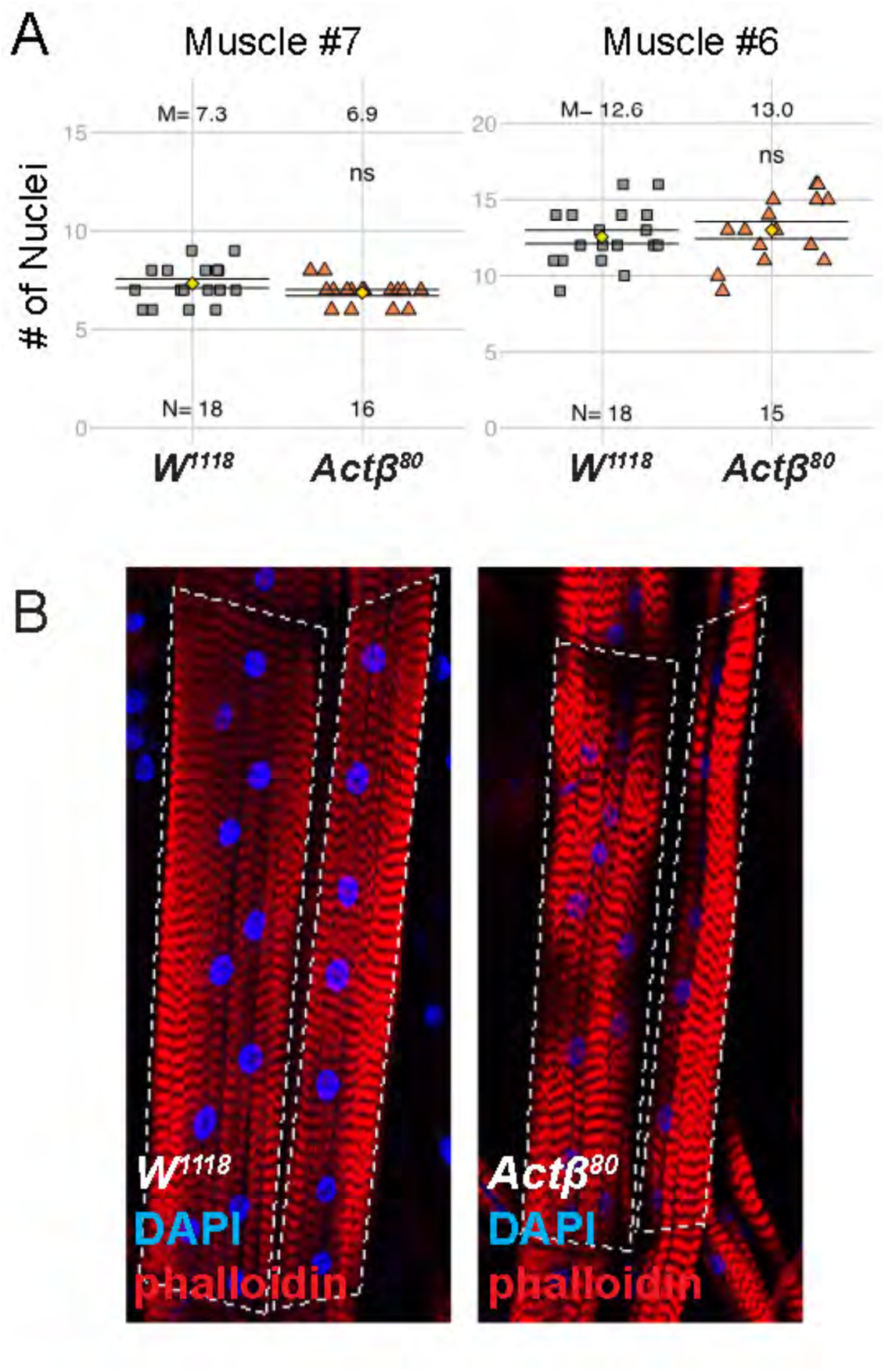
*Actβ* does not affect muscle nuclei number. 3rd instar larvae were filleted and stained with DAPI (blue) and phalloidin (red). (A) Quantification of the number of nuclei in muscle #6 and #7 for *w^1118^* controls and *Actβ^ed80^*mutants shows no significant difference. (B) Representative image of muscle fiber (red) with stained nuclei (blue) for *w^1118^* controls and *Actβ^ed80^*mutant. Each image has muscle #6 (left) and muscle #7 (right) outlined with dashed line. N is shown for each group in A, ns = not significant

**Figure S2.**
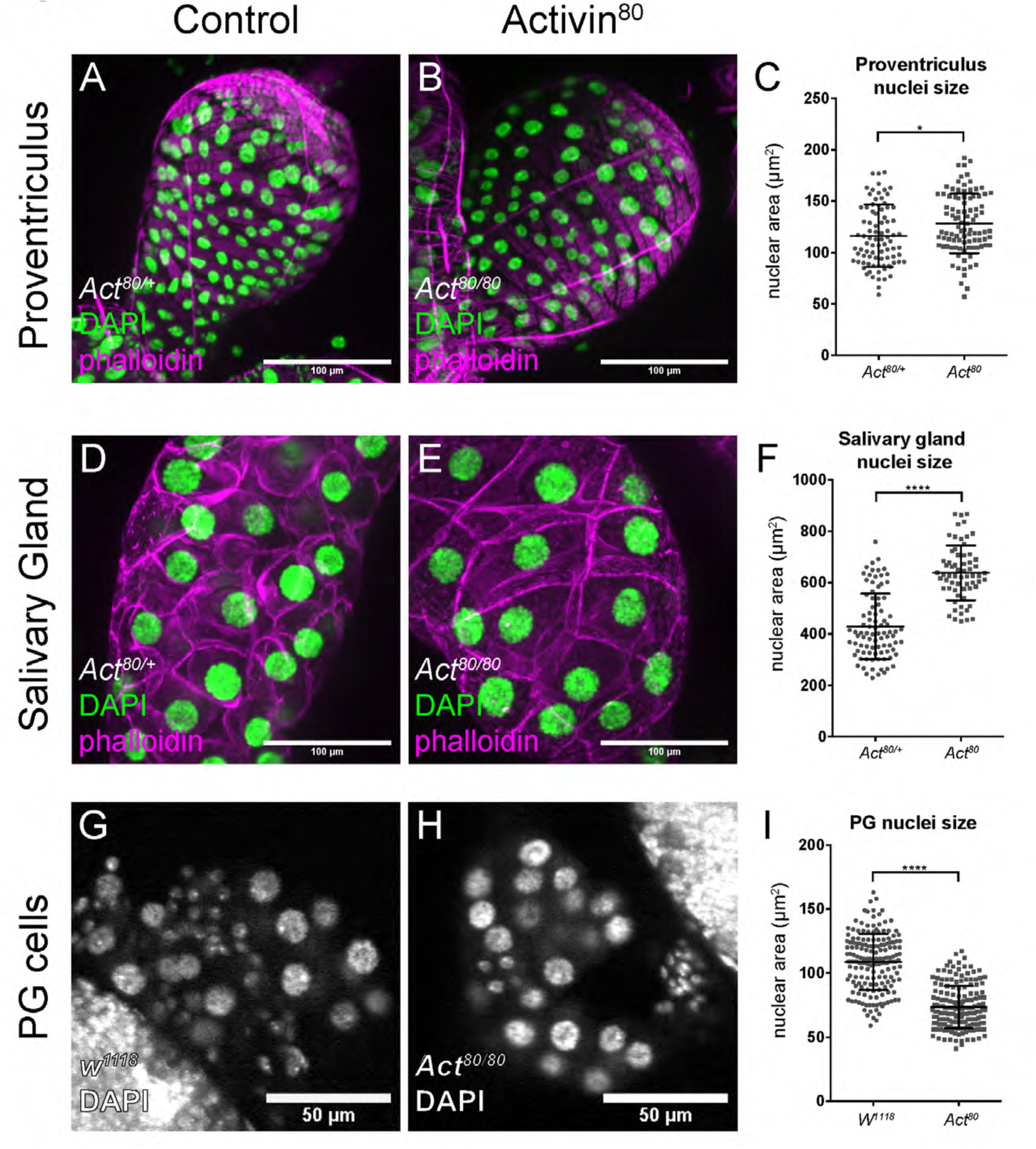
Size of various polyploid nuclei in *Actβ* mutants. Proventriculus, salivary gland, and the prothoracic gland of 3rd instar wandering larvae were dissected, stained with DAPI (green, A, B, D, E; grey, G, H) and phalloidin (magenta, A, B, D, E) to quantify nuclei size. (A-C) The nuclei of the cells of the proventriculus in *Actβ^ed80^*mutants are 10% larger. (D-F) The nuclei of the salivary gland cells in *Actβ^ed80^*mutants are 48% larger. (G-I) The nuclei of the prothoracic gland cells are −32% smaller. Size of scale bar indicated on the image. For each group n = 60-170.

**Figure S3.**
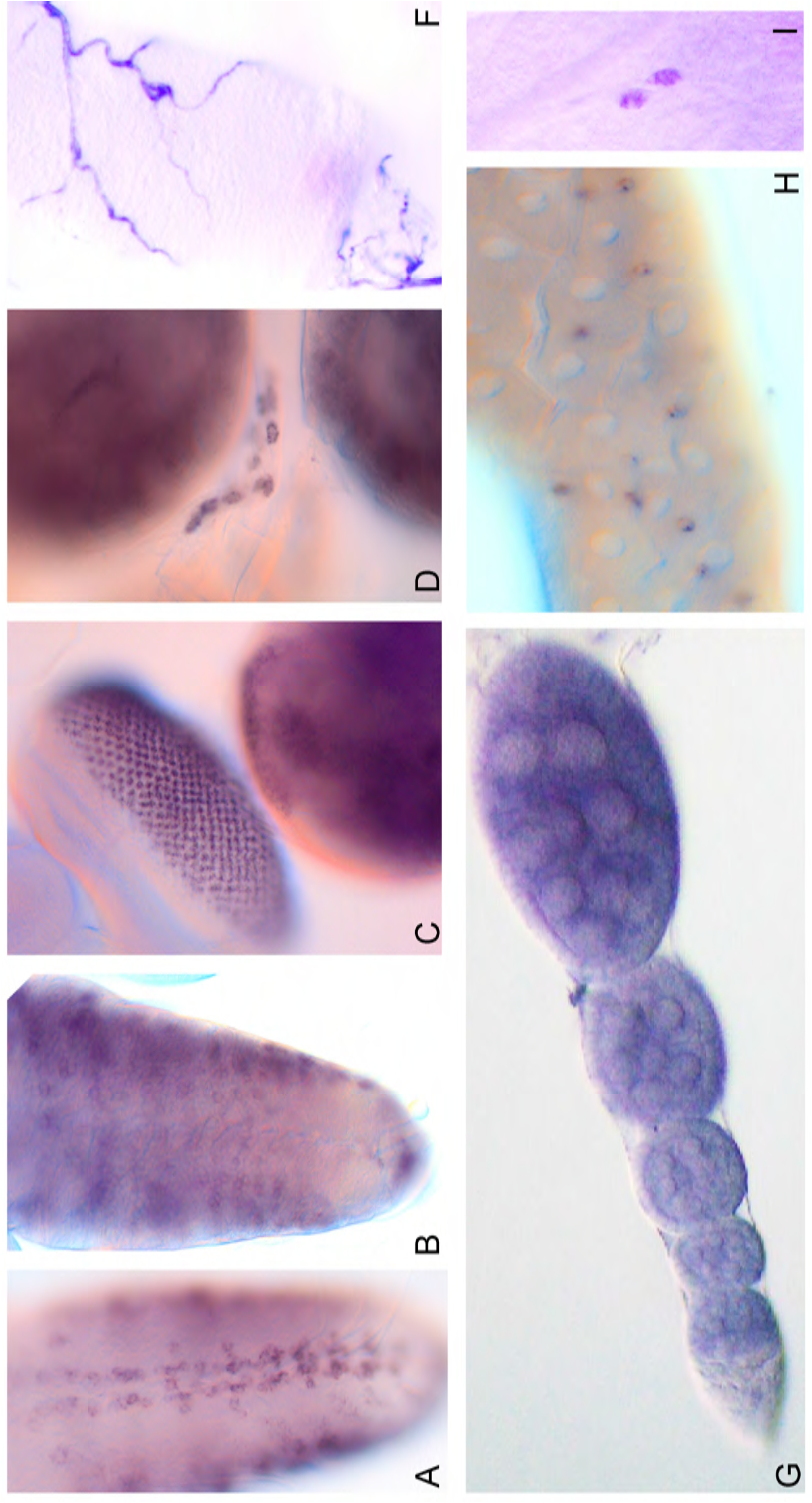
RNA *in situ* hybridization of *Actβ* shows expression in various tissues. (A-I) *Actβ* is expressed in cell body of motoneurons (A) and neuroendocrine cells (B) in the ventral ganglion of 3rd instar larval CNS, (C) eye disc, (D) CC cells, (F) trachea, (G) maturing follicle cells, (H) enteroendocrine cells in the midgut, (I) two PNS neurons.

**Figure S4.**
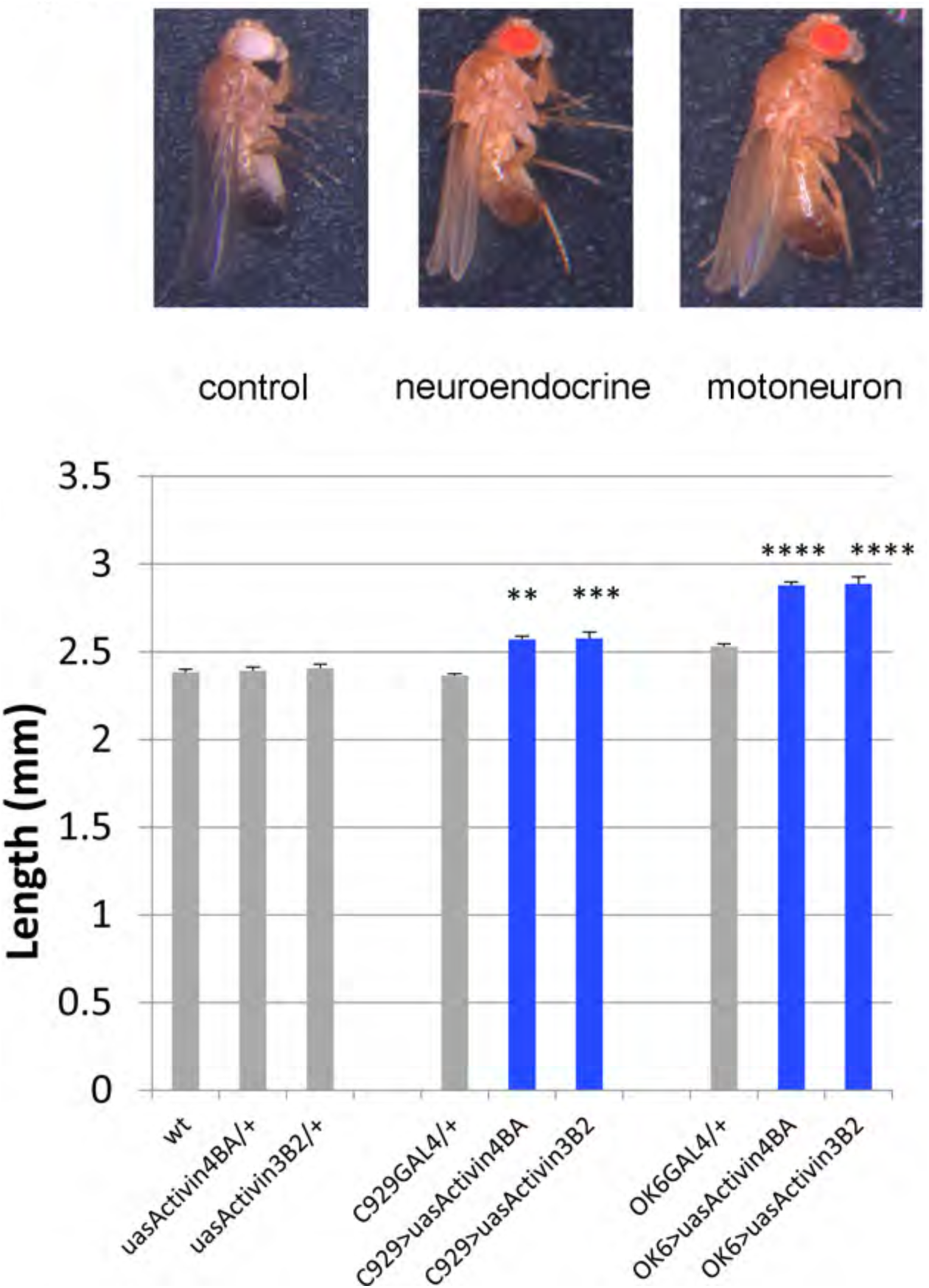
Overexpression of *Actβ* in motoneurons or neuroendocrine cells increases adult body size. Relative to controls (UAS-*Actβ*, left) overexpression of *Actβ* (line 3B2 or 4BA) in motoneurons (OK6-Gal4, right) or neuroendocrine cell (C929-Gal4, middle) results in larger animals. The body length for the indicated genotype shown (bottom panel) with representative images of adult flies (top panel). n = 20-30 individuals, bars indicate mean ± SEM.

**Figure S5.**
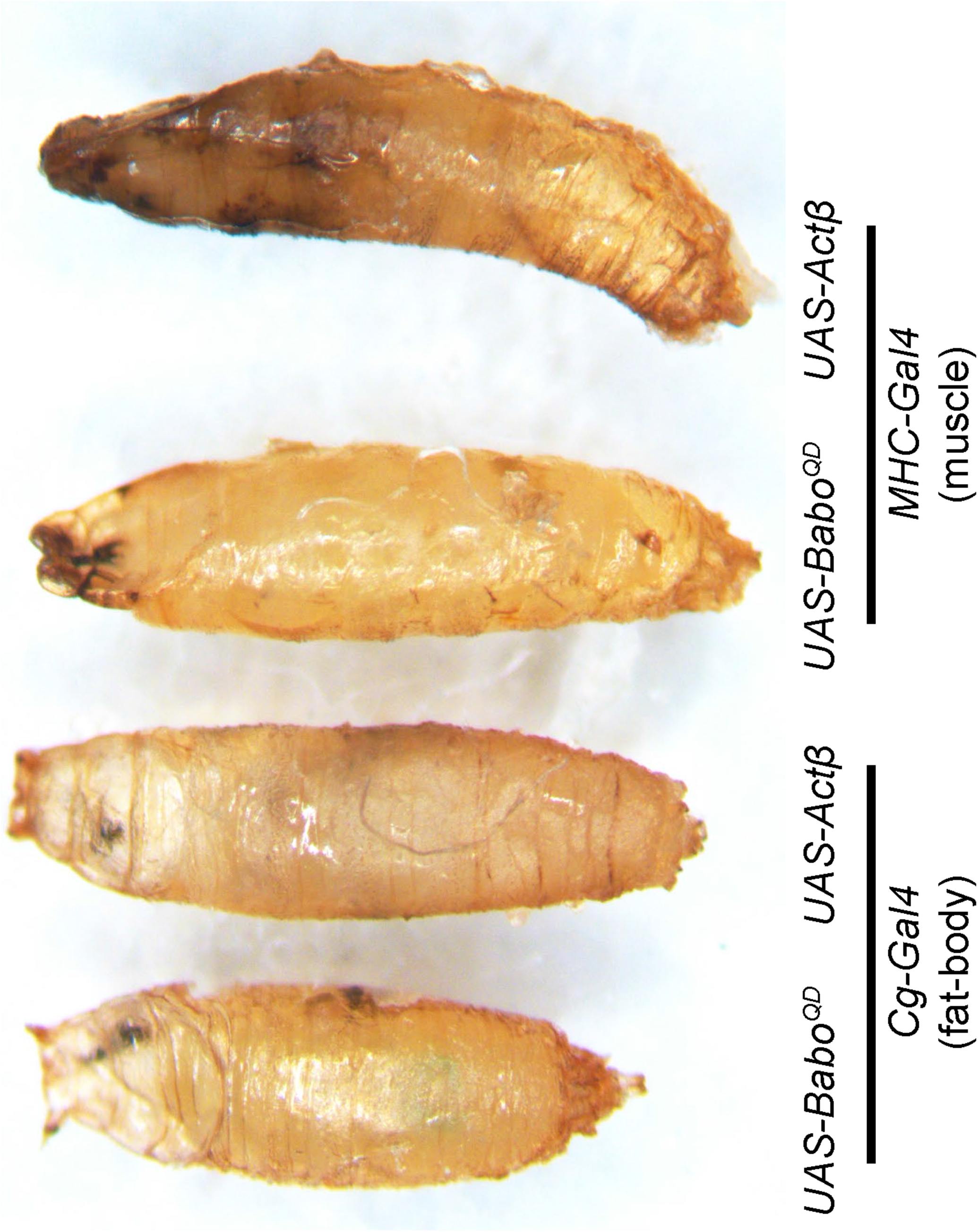
Hyperactivation of Activin signaling in muscles but not fatbody is larval lethal. Images of pupae resulting from overexpression of constitutively activated Babo (Babo^QD^) or *Actβ* in the fat-body (*Cg-Gal4,* left) or muscles (*MHC-Gal4*, right). Note that the left most pupa in which activated Babo was expressed in the fatbody produces a normal pupa that gives rise to a viable adult. However expression of the ligand (Actβ) in the fatbody produces a larval lethal phenotype similar to that produced by either overexpression of Actβ or activated receptor in muscle. We conclude that overexpression of the ligand in the fatbody produces a lethal phenotype by non-autonomous activity in the muscles.

**Figure S6.**
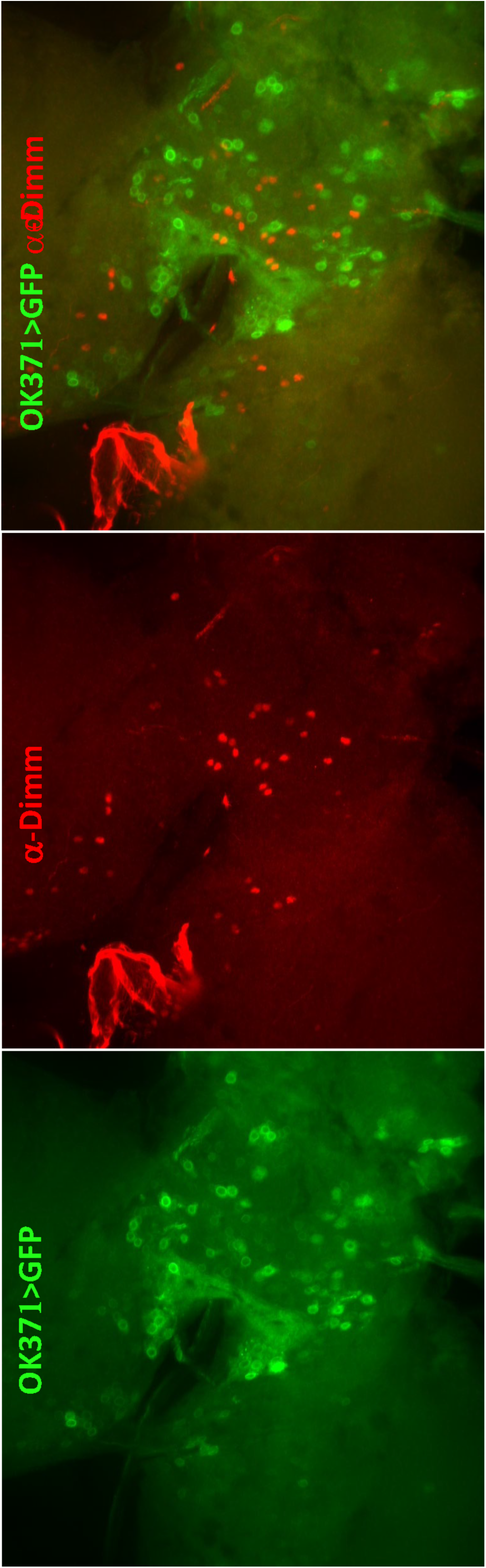
OK372-Gal4 is not expressed in Dimm + neuroendocrine cells. Third instar larvae brains of *OK371-Gal4>UAS-GFP* (green) were dissected and stained with α-Dimm (red). We find no overlap in signals demonstrating that OK371-Gal4 is not expressed in Dimm+ cells (merge panel, right).

**Figure S7.**
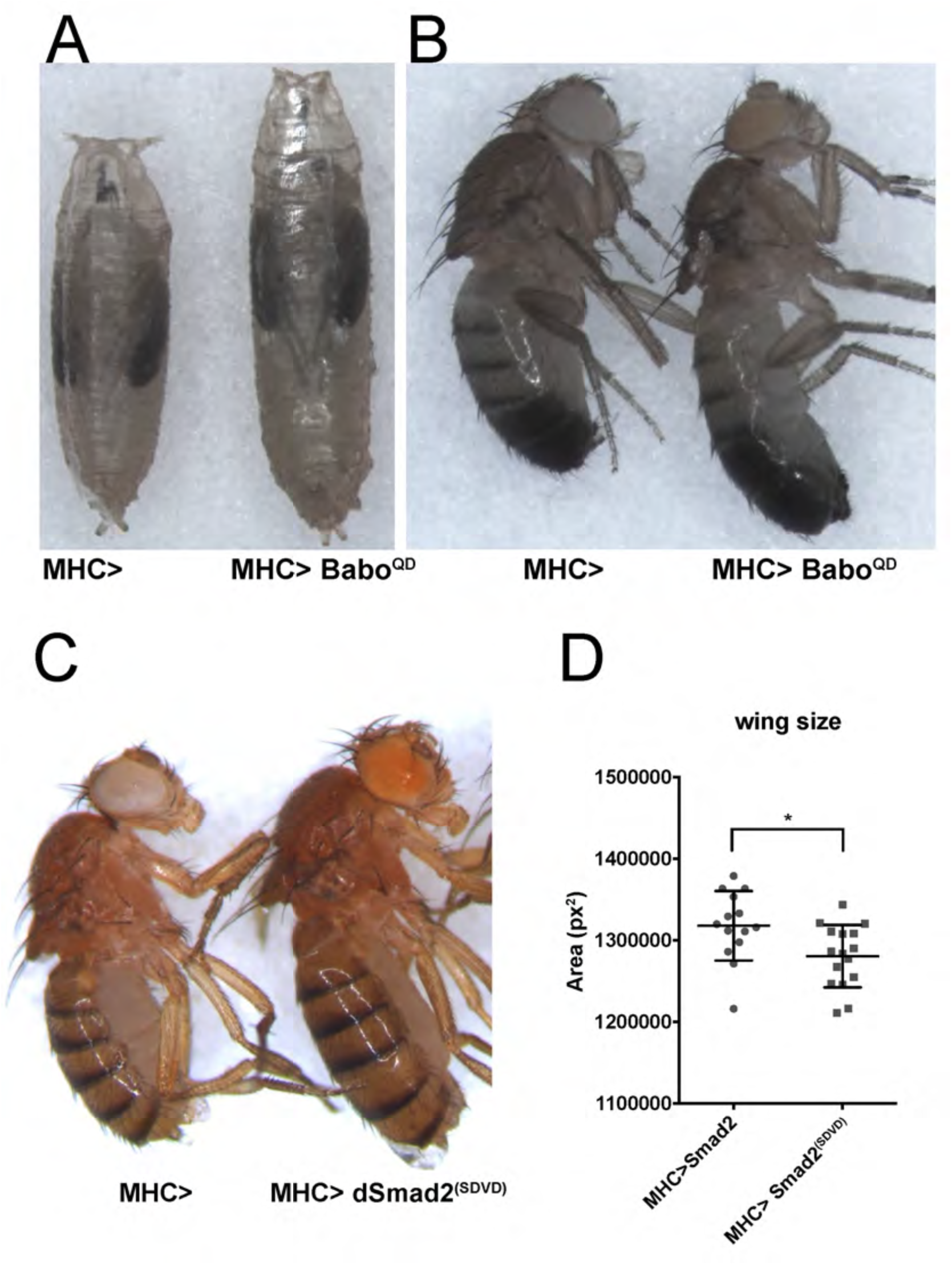
Muscle specific activation of Activin signaling increases body size but not wing size. (A-B) Overexpression of activated Babo (Babo^QD^) in muscles results in larger pupae (A) and adult larger adult flies (B). (C-D) Overexpression of activated dSmad2 (dSmad2^SDVD^) in muscles also results in larger adult flies. (D) adult wings of *MHC>dSmad2^SDVD^* are smaller than *MHC-Gal4* control.

## References

Augustin, H., K. McGourty, J. R. Steinert, H. M. Cocheme, J. Adcott et al., 2017 Myostatin-like proteins regulate synaptic function and neuronal morphology. Development 144: 2445–2455.

Bohni, R., J. Riesgo-Escovar, S. Oldham, W. Brogiolo, H. Stocker et al., 1999 Autonomous control of cell and organ size by CHICO, a Drosophila homolog of vertebrate IRS1-4. Cell 97: 865–875.

Boulan, L., M. Milan and P. Leopold, 2015 The Systemic Control of Growth. Cold Spring Harb Perspect Biol 7.

Brogiolo, W., H. Stocker, T. Ikeya, F. Rintelen, R. Fernandez et al., 2001 An evolutionarily conserved function of the Drosophila insulin receptor and insulin-like peptides in growth control. Curr Biol 11: 213–221.

Chen, C., J. Jack and R. S. Garofalo, 1996 The Drosophila insulin receptor is required for normal growth. Endocrinology 137: 846–856.

Chng, W. A., M. S. B. Sleiman, F. Schupfer and B. Lemaitre, 2014 Transforming growth factor beta/activin signaling functions as a sugar-sensing feedback loop to regulate digestive enzyme expression. Cell Rep 9: 336–348.

Church, R. B., and F. W. Robertson, 1966 A biochemical study of growth of Drsophila melanogaster J. Exp. Zool 162: 337–351.

De Loof, A., T. Vandersmissen, E. Marchal and L. Schoofs, 2015 Initiation of metamorphosis and control of ecdysteroid biosynthesis in insects: The interplay of absence of Juvenile hormone, PTTH, and Ca(2+)-homeostasis. Peptides 68: 120–129.

Demontis, F., and N. Perrimon, 2009 Integration of Insulin receptor/Foxo signaling and dMyc activity during muscle growth regulates body size in Drosophila. Development 136: 983–993.

Derynck, R., W. M. Gelbart, R. M. Harland, C. H. Heldin, S. E. Kern et al., 1996 Nomenclature: vertebrate mediators of TGFbeta family signals. Cell 87: 173.

Droujinine, I. A., and N. Perrimon, 2016 Interorgan Communication Pathways in Physiology: Focus on Drosophila. Annu Rev Genet 50: 539–570.

Elashry, M. I., A. Otto, A. Matsakas, S. E. El-Morsy and K. Patel, 2009 Morphology and myofiber composition of skeletal musculature of the forelimb in young and aged wild type and myostatin null mice. Rejuvenation Res 12: 269–281.

Elkasrawy, M. N., and M. W. Hamrick, 2010 Myostatin (GDF-8) as a key factor linking muscle mass and bone structure. J Musculoskelet Neuronal Interact 10: 56–63.

Gesualdi, S. C., and T. E. Haerry, 2007 Distinct signaling of Drosophila Activin/TGF-beta family members. Fly (Austin) 1: 212–221.

Ghosh, A. C., and M. B. O’Connor, 2014 Systemic Activin signaling independently regulates sugar homeostasis, cellular metabolism, and pH balance in Drosophila melanogaster. Proc Natl Acad Sci U S A 111: 5729–5734.

Goberdhan, D. C., N. Paricio, E. C. Goodman, M. Mlodzik and C. Wilson, 1999 Drosophila tumor suppressor PTEN controls cell size and number by antagonizing the Chico/PI3-kinase signaling pathway. Genes Dev 13: 3244–3258.

Gokhale, R. H., T. Hayashi, C. D. Mirque and A. W. Shingleton, 2016 Intra-organ growth coordination in Drosophila is mediated by systemic ecdysone signaling. Dev Biol 418: 135–145.

Hariharan, I. K., D. B. Wake and M. H. Wake, 2015 Indeterminate Growth: Could It Represent the Ancestral Condition? Cold Spring Harb Perspect Biol 8: a019174.

Hata, A., and Y. G. Chen, 2016 TGF-beta Signaling from Receptors to Smads. Cold Spring Harb Perspect Biol 8.

Heldin, C. H., and A. Moustakas, 2016 Signaling Receptors for TGF-beta Family Members. Cold Spring Harb Perspect Biol 8.

Hill, C. S., 2016 Transcriptional Control by the SMADs. Cold Spring Harb Perspect Biol 8.

Hirschhorn, J. N., and G. Lettre, 2009 Progress in genome-wide association studies of human height. Horm Res 71 Suppl 2: 5–13.

Jensen, P. A., X. Zheng, T. Lee and M. B. O’Connor, 2009 The Drosophila Activin-like ligand Dawdle signals preferentially through one isoform of the Type-I receptor Baboon. Mech Dev 126: 950–957.

Kahlem, P., and S. J. Newfeld, 2009 Informatics approaches to understanding TGFbeta pathway regulation. Development 136: 3729–3740.

Kim, M. J., and M. B. O’Connor, 2014 Anterograde Activin signaling regulates postsynaptic membrane potential and GluRIIA/B abundance at the Drosophila neuromuscular junction. PLoS One 9: e107443.

Koyama, T., and C. K. Mirth, 2018 Unravelling the diversity of mechanisms through which nutrition regulates body size in insects. Curr Opin Insect Sci 25: 1–8.

Kronenberg, H. M., 2003 Developmental regulation of the growth plate. Nature 423: 332–336.

Lamouille, S., and R. Derynck, 2007 Cell size and invasion in TGF-beta-induced epithelial to mesenchymal transition is regulated by activation of the mTOR pathway. J Cell Biol 178: 437–451.

Lee, J. Y., N. S. Hopkinson and P. R. Kemp, 2011 Myostatin induces autophagy in skeletal muscle in vitro. Biochem Biophys Res Commun 415: 632–636.

Leevers, S. J., D. Weinkove, L. K. MacDougall, E. Hafen and M. D. Waterfield, 1996 The Drosophila phosphoinositide 3-kinase Dp110 promotes cell growth. EMBO J 15: 6584–6594.

Lokireddy, S., I. W. Wijesoma, S. K. Sze, C. McFarlane, R. Kambadur et al., 2012 Identification of atrogin-1-targeted proteins during the myostatin-induced skeletal muscle wasting. Am J Physiol Cell Physiol 303: C512–529.

Macias, M. J., P. Martin-Malpartida and J. Massague, 2015 Structural determinants of Smad function in TGF-beta signaling. Trends Biochem Sci 40: 296–308.

Madaan, U., E. Yzeiraj, M. Meade, J. F. Clark, C. A. Rushlow et al., 2018 BMP Signaling Determines Body Size via Transcriptional Regulation of Collagen Genes in Caenorhabditis elegans. Genetics 210: 1355–1367.

Makhijani, K., B. Alexander, D. Rao, S. Petraki, L. Herboso et al., 2017 Regulation of Drosophila hematopoietic sites by Activin-beta from active sensory neurons. Nat Commun 8: 15990.

Marques, G., H. Bao, T. E. Haerry, M. J. Shimell, P. Duchek et al., 2002 The Drosophila BMP type II receptor Wishful Thinking regulates neuromuscular synapse morphology and function. Neuron 33: 529–543.

Matsakas, A., A. Otto, M. I. Elashry, S. C. Brown and K. Patel, 2010 Altered primary and secondary myogenesis in the myostatin-null mouse. Rejuvenation Res 13: 717–727.

McBrayer, Z., H. Ono, M. Shimell, J. P. Parvy, R. B. Beckstead et al., 2007 Prothoracicotropic hormone regulates developmental timing and body size in Drosophila. Dev Cell 13: 857–871.

McNabb, S. L., J. D. Baker, J. Agapite, H. Steller, L. M. Riddiford et al., 1997 Disruption of a behavioral sequence by targeted death of peptidergic neurons in Drosophila. Neuron 19: 813–823.

McPherron, A. C., A. M. Lawler and S. J. Lee, 1997 Regulation of skeletal muscle mass in mice by a new TGF-beta superfamily member. Nature 387: 83–90.

McPherron, A. C., and S. J. Lee, 1997 Double muscling in cattle due to mutations in the myostatin gene. Proc Natl Acad Sci U S A 94: 12457–12461.

Mirth, C. K., and A. W. Shingleton, 2012 Integrating body and organ size in Drosophila: recent advances and outstanding problems. Front Endocrinol (Lausanne) 3: 49.

Mirth, C. K., and A. W. Shingleton, 2014 The roles of juvenile hormone, insulin/target of rapamycin, and ecydsone signaling in regulating body size in Drosophila. Commun Integr Biol 7.

Mirth, C. K., H. Y. Tang, S. C. Makohon-Moore, S. Salhadar, R. H. Gokhale et al., 2014 Juvenile hormone regulates body size and perturbs insulin signaling in Drosophila. Proc Natl Acad Sci U S A 111: 7018–7023.

Nijhout, H. F., and V. Callier, 2015 Developmental mechanisms of body size and wing-body scaling in insects. Annu Rev Entomol 60: 141–156.

Nijhout, H. F., and L. W. Grunert, 2010 The cellular and physiological mechanism of wing-body scaling in Manduca sexta. Science 330: 1693–1695.

Oldham, S., J. Montagne, T. Radimerski, G. Thomas and E. Hafen, 2000 Genetic and biochemical characterization of dTOR, the Drosophila homolog of the target of rapamycin. Genes Dev 14: 2689–2694.

Park, D., J. A. Veenstra, J. H. Park and P. H. Taghert, 2008 Mapping peptidergic cells in Drosophila: where DIMM fits in. PLoS One 3: e1896.

Parker, N. F., and A. W. Shingleton, 2011 The coordination of growth among Drosophila organs in response to localized growth-perturbation. Dev Biol 357: 318–325.

Prokop, A., 2006 Organization of the efferent system and structure of neuromuscular junctions in Drosophila. Int Rev Neurobiol 75: 71–90.

Prokop, A., and I. A. Meinertzhagen, 2006 Development and structure of synaptic contacts in Drosophila. Semin Cell Dev Biol 17: 20–30.

Ren, X., J. Sun, B. E. Housden, Y. Hu, C. Roesel et al., 2013 Optimized gene editing technology for Drosophila melanogaster using germ line-specific Cas9. Proc Natl Acad Sci U S A 110: 19012–19017.

Rewitz, K. F., N. Yamanaka, L. I. Gilbert and M. B. O’Connor, 2009 The insect neuropeptide PTTH activates receptor tyrosine kinase torso to initiate metamorphosis. Science 326: 1403–1405.

Rewitz, K. F., N. Yamanaka and M. B. O’Connor, 2013 Developmental checkpoints and feedback circuits time insect maturation. Curr Top Dev Biol 103: 1–33.

Riddiford, L. M., J. W. Truman, C. K. Mirth and Y. C. Shen, 2010 A role for juvenile hormone in the prepupal development of Drosophila melanogaster. Development 137: 1117–1126.

Rulifson, E. J., S. K. Kim and R. Nusse, 2002 Ablation of insulin-producing neurons in flies: growth and diabetic phenotypes. Science 296: 1118–1120.

Sartori, R., E. Schirwis, B. Blaauw, S. Bortolanza, J. Zhao et al., 2013 BMP signaling controls muscle mass. Nat Genet 45: 1309–1318.

Schindelin, J., I. Arganda-Carreras, E. Frise, V. Kaynig, M. Longair et al., 2012 Fiji: an open-source platform for biological-image analysis. Nat Methods 9: 676–682.

Sebo, Z. L., H. B. Lee, Y. Peng and Y. Guo, 2014 A simplified and efficient germline-specific CRISPR/Cas9 system for Drosophila genomic engineering. Fly (Austin) 8: 52–57.

Shim, K. S., 2015 Pubertal growth and epiphyseal fusion. Ann Pediatr Endocrinol Metab 20: 8–12.

Shimell, M., X. Pan, F. A. Martin, A. C. Ghosh, P. Leopold et al., 2018 Prothoracicotropic hormone modulates environmental adaptive plasticity through the control of developmental timing. Development 145.

Shingleton, A. W., C. M. Estep, M. V. Driscoll and I. Dworkin, 2009 Many ways to be small: different environmental regulators of size generate distinct scaling relationships in Drosophila melanogaster. Proc Biol Sci 276: 2625–2633.

Shingleton, A. W., and W. A. Frankino, 2013 New perspectives on the evolution of exaggerated traits. Bioessays 35: 100–107.

Shingleton, A. W., W. A. Frankino, T. Flatt, H. F. Nijhout and D. J. Emlen, 2007 Size and shape: the developmental regulation of static allometry in insects. Bioessays 29: 536–548.

Siegmund, T., and G. Korge, 2001 Innervation of the ring gland of Drosophila melanogaster. J Comp Neurol 431: 481–491.

Simpson, P., P. Berreur and J. Berreur-Bonnenfant, 1980 The initiation of pupariation in Drosophila: dependence on growth of the imaginal discs. J Embryol Exp Morphol 57: 155–165.

Simpson, P., and H. A. Schneiderman, 1975 Isolation of temperature sensitive mutations blocking clone development inDrosophila melanogaster, and the effects of a temperature sensitive cell lethal mutation on pattern formation in imaginal discs. Wilehm Roux Arch Dev Biol 178: 247–275.

Song, W., D. Cheng, S. Hong, B. Sappe, Y. Hu et al., 2017a Midgut-Derived Activin Regulates Glucagon-like Action in the Fat Body and Glycemic Control. Cell Metab 25: 386–399.

Song, W., E. Owusu-Ansah, Y. Hu, D. Cheng, X. Ni et al., 2017b Activin signaling mediates muscle-to-adipose communication in a mitochondria dysfunction-associated obesity model. Proc Natl Acad Sci U S A.

Stieper, B. C., M. Kupershtok, M. V. Driscoll and A. W. Shingleton, 2008 Imaginal discs regulate developmental timing in Drosophila melanogaster. Dev Biol 321: 18–26.

Stocker, H., T. Radimerski, B. Schindelholz, F. Wittwer, P. Belawat et al., 2003 Rheb is an essential regulator of S6K in controlling cell growth in Drosophila. Nat Cell Biol 5: 559–565.

Ting, C. Y., T. Herman, S. Yonekura, S. Gao, J. Wang et al., 2007 Tiling of r7 axons in the Drosophila visual system is mediated both by transduction of an activin signal to the nucleus and by mutual repulsion. Neuron 56: 793–806.

Ting, C. Y., P. G. McQueen, N. Pandya, T. Y. Lin, M. Yang et al., 2014 Photoreceptor-derived activin promotes dendritic termination and restricts the receptive fields of first-order interneurons in Drosophila. Neuron 81: 830–846.

Trendelenburg, A. U., A. Meyer, D. Rohner, J. Boyle, S. Hatakeyama et al., 2009 Myostatin reduces Akt/TORC1/p70S6K signaling, inhibiting myoblast differentiation and myotube size. Am J Physiol Cell Physiol 296: C1258–1270.

Tuck, S., 2014 The control of cell growth and body size in Caenorhabditis elegans. Exp Cell Res 321: 71–76.

Upadhyay, A., L. Moss-Taylor, M. J. Kim, A. C. Ghosh and M. B. O’Connor, 2017 TGF-beta Family Signaling in Drosophila. Cold Spring Harb Perspect Biol 9.

Winbanks, C. E., J. L. Chen, H. Qian, Y. Liu, B. C. Bernardo et al., 2013 The bone morphogenetic protein axis is a positive regulator of skeletal muscle mass. J Cell Biol 203: 345–357.

Wood, A. R., T. Esko, J. Yang, S. Vedantam, T. H. Pers et al., 2014 Defining the role of common variation in the genomic and biological architecture of adult human height. Nat Genet 46: 1173–1186.

Wu, Q., T. Wen, G. Lee, J. H. Park, H. N. Cai et al., 2003 Developmental control of foraging and social behavior by the Drosophila neuropeptide Y-like system. Neuron 39: 147–161.

Wu, Q., Y. Zhang, J. Xu and P. Shen, 2005 Regulation of hunger-driven behaviors by neural ribosomal S6 kinase in Drosophila. Proc Natl Acad Sci U S A 102: 13289–13294.

Yamanaka, N., K. F. Rewitz and M. B. O’Connor, 2013a Ecdysone control of developmental transitions: lessons from Drosophila research. Annu Rev Entomol 58: 497–516.

Yamanaka, N., N. M. Romero, F. A. Martin, K. F. Rewitz, M. Sun et al., 2013b Neuroendocrine control of Drosophila larval light preference. Science 341: 1113–1116.

Zhou, X., J. L. Wang, J. Lu, Y. Song, K. S. Kwak et al., 2010 Reversal of cancer cachexia and muscle wasting by ActRIIB antagonism leads to prolonged survival. Cell 142: 531–543.

Zhu, C. C., J. Q. Boone, P. A. Jensen, S. Hanna, L. Podemski et al., 2008 Drosophila Activin- and the Activin-like product Dawdle function redundantly to regulate proliferation in the larval brain. Development 135: 513–521.

